# High sorbic acid resistance of *Penicillium roqueforti* is mediated by the SORBUS gene cluster

**DOI:** 10.1101/2022.02.10.479849

**Authors:** Maarten Punt, Sjoerd J. Seekles, Jisca L. van Dam, Connor de Adelhart Toorop, Raithel R Martina, Jos Houbraken, Arthur F. J. Ram, Han A. B. Wösten, Robin A. Ohm

## Abstract

*Penicillium roqueforti* is a major food-spoilage fungus known for its high resistance to the food preservative sorbic acid. Here, we demonstrate that the minimum inhibitory concentration of undissociated sorbic acid (MIC_u_) ranges between 4.2 and 21.2 mM when 34 *P. roqueforti* strains were grown on malt extract broth. A genome-wide association study revealed that the six most resistant strains contained the 180 kbp gene cluster SORBUS, which was absent in the other 28 strains. In addition, a SNP analysis revealed five genes outside the SORBUS cluster that may be linked to sorbic acid resistance. A partial SORBUS knock-out (>100 of 180 kbp) in a resistant strain reduced sorbic acid resistance to similar levels as observed in the sensitive strains. Whole genome transcriptome analysis revealed a small set of genes present in both resistant and sensitive *P. roqueforti* strains that were differentially expressed in the presence of the weak acid. These genes could explain why *P. roqueforti* is more resistant to sorbic acid when compared to other fungi, even in the absence of the SORBUS cluster. Together, the MIC_u_ of 21.2 mM makes *P. roqueforti* among the most sorbic acid-resistant fungi, if not the most resistant fungus, which is mediated by the SORBUS gene cluster.

**Author summary:** Chemical preservatives, such as sorbic acid, are often used in food to prevent spoilage by fungi, yet some fungi are particularly well-suited to deal with these preservatives. First, we investigated the resistance of 34 *Penicillium roqueforti* strains to various food preservatives. This revealed that some strains were highly resistant to sorbic acid, while others are more sensitive. Next, we used DNA sequencing to compare the genetic variation between these strains and discovered a specific genetic region (SORBUS) that is unique to the resistant strains. Through comparative analysis with other fungal species the SORBUS region was studied in more detail and with the use of genetic engineering tools we removed this unique region. Finally, the mutant lacking the SORBUS region was confirmed to have lost its sorbic acid resistance. This finding is of particular interest as it suggests that only some, not all, *P. roqueforti* strains are potent spoilers and that specific genetic markers could help in the identification of resistant strains.

## Introduction

Fungi are responsible for 5–10 % of all food spoilage [1] resulting in the production of off-flavors, discoloration and acidification [1,2]. Some species also produce mycotoxins, such as PR toxin and roquefortine C in the case of *Penicillium roqueforti* [3]. Toxins partially contribute to the high incidence of food-borne diseases affecting up to 30 % of the people in industrialized countries every year [4].

Food spoilage by filamentous fungi is believed to be mainly caused by spores that are spread through the air, water or other vectors like insects [5]. Preservation techniques such as pasteurization, fermentation, cooling or the addition of preservatives are used to reduce spoilage [6]. Some of the most applied preservatives are weak organic acids such as benzoic, propionic and sorbic acid. *Paecilomyces variotii, Penicillium paneum, Penicillium carneum* and *P. roqueforti* are among the few filamentous fungi capable of spoiling products containing weak acids, and are therefore called preservative-resistant moulds [7]. Weak-acid preservatives inhibit microbial growth, but their mode-of-action is not completely understood. According to the classical ‘weak-acid preservative theory’ the antimicrobial activity of weak acids is derived from their undissociated form that can pass the plasma membrane. These weak acids dissociate in the cytosol due to its neutral pH, and inhibit growth through acidification of the cytoplasm [8]. The inhibitory activity of sorbic acid at pH 6.5 and the correlation of sorbic acid resistance with ethanol tolerance in *Saccharomyces cerevisiae* suggest that this weak acid can also act as a membrane-active compound [8].

In *A. niger*, sorbic acid resistance is mediated by the phenylacrylic acid decarboxylase gene *padA*, and the putative 4-hydroxybenzoate decarboxylase gene, known as the cinnamic acid decarboxylase (*cdcA*) gene [9,10]. These genes encode proteins that catalyze the conversion of sorbic acid into 1,3-pentadiene. Genes *padA* and *cdcA* are regulated by the sorbic acid decarboxylase regulator (SdrA), which is a Zn2Cys6-finger transcription factor [10]. These three genes are present on the same genetic locus in *A. niger* with *sdrA* being flanked by *cdcA* and *padA*. Orthologs of *cdcA* and *padA* are also clustered in *S. cerevisiae*, but this is not the case for the *sdrA* homologue [10,11]. Inactivation of *cdcA, padA* or *sdrA* results in reduced growth of *A. niger* on sorbic acid and cinnamic acid [9]. The fact that growth is not abolished and the fact that *padA* transcript levels are less affected than that of *cdcA* on sorbic acid upon deletion of *sdrA* suggests that an additional regulator is involved [9]. Indeed, the weak-acid regulator WarA is also involved in sorbic acid resistance as well as other weak acids such as propionic and benzoic acid [12].

Weak-acid resistance in fungi is subject to intra- and inter-strain heterogeneity. Intra-strain heterogeneity has been observed in *Zygosaccharomyces bailii* with the existence of a small sub-population of cells that are more resistant (MIC = 7.6 mM) to sorbic acid than the sensitive population (MIC = 3 mM) [13], while inter-strain resistance has been observed in *P. roqueforti* with the existence of sorbic acid-resistant and sorbic acid-sensitive strains [14]. Genome analysis of *P. roqueforti* strains have yielded clues about adaptive divergence in this species. Genomic islands with high identity are present in distant *Penicillium* species, while they are absent in closely related species, supporting the hypothesis of recent horizontal gene transfer events [15,16]. For instance, the presence of two large genomic regions, *Wallaby* and *CheesyTer* correlates with faster growth and functions relevant in a cheese matrix, respectively [16,17]. Noteworthy, the *CheesyTer* and *Wallaby* regions are only present in cheese isolates other than Roquefort, indicating that the Roquefort isolate population is potentially more closely related to the ‘ancestral’ *P. roqueforti* populations [18].

In this study, sorbic acid resistance of 34 *P. roqueforti* strains was assessed. A genome-wide association analysis revealed the presence of the 180 kbp gene cluster SORBUS in the six most resistant strains. A partial SORBUS knockout in such a strain showed a reduced sorbic acid resistance similar to that of the other 28 sensitive *P. roqueforti* strains.

## Results

### Weak-acid sensitivity screening

Weak-acid sensitivity of 34 *P. roqueforti* wild-type strains was assessed on MEA plates supplemented with 5 mM propionic, sorbic or benzoic acid, which corresponds to 4.42, 4.25 and 3.07 mM undissociated acid, respectively. Three strains had been isolated from blue-veined cheeses such as Roquefort and the other 31 strains had been isolated from non-cheese environments (mostly related to spoiled food) (Table S1). The colony surface area was determined after five days of growth (Fig 1). The inhibitory effect of propionic acid at the tested concentration was limited for most strains, since 26 out of 34 strains grew to > 80 % of the colony surface area reached under control conditions. Strains DTO012A1 and DTO012A8 even showed an increased colony size (up to 120 %) when compared to the control. The inhibitory effects of sorbic and benzoic acid were more pronounced. MEA supplemented with potassium sorbate reduced colony area for all 34 *P. roqueforti* strains. The surface area under sorbic acid stress ranged from 0 to 80 % of the surface area reached under control conditions. Strains DTO006G1, DTO006G7, DTO013E5 and DTO013F2 showed the highest sorbic acid resistance, followed by DTO046C5. Benzoic acid was the most inhibitory compound, resulting in a maximum colony surface area between 0 and 20 % of the control. The most benzoic acid-resistant strains were DTO013F2 and DTO013E5. As these strains were also among the most sorbic acid-resistant strains, similar resistance mechanisms may be involved to cope with benzoic and sorbic acid stress.

**Fig 1.**
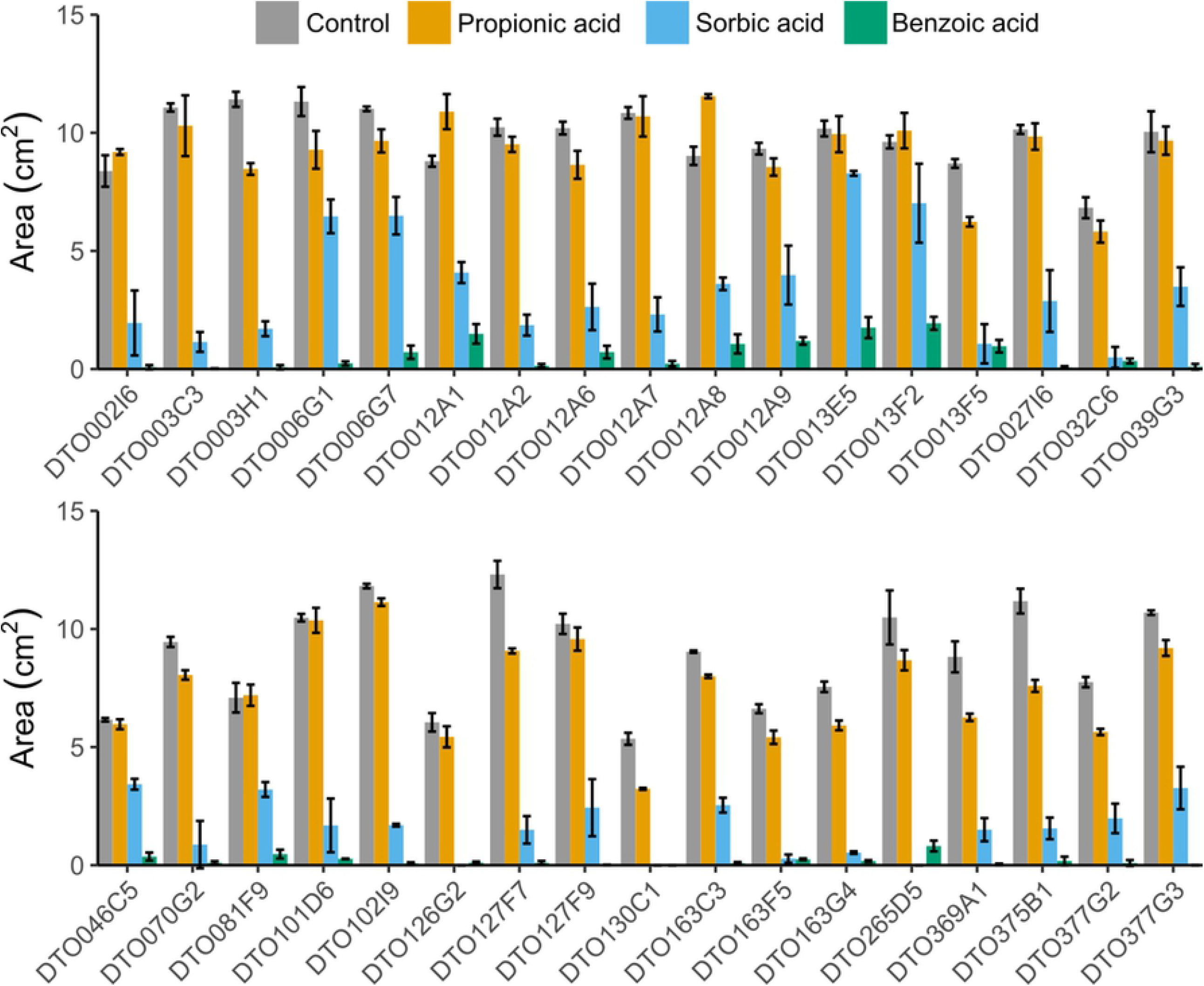
Weak-acid screening shows variabe resistances to multiple acids. Average colony size (cm^2^) of 34 *P. roqueforti* strains after five days of growth on MEA (pH 4.0) (grey) or MEA (pH 4.0) supplemented with 5 mM propionic acid (orange), sorbic acid (blue) or benzoic acid (green). Strains DTO006G1, DTO006G7, DTO013E5 and DTO013F2 are relatively resistant to sorbic acid. Error bars indicate standard deviation of biologically independent replicates.

Sorbic acid resistance was further analyzed by determining the MIC_u_ values of sorbic acid for the 34 *P. roqueforti* strains. The four strains with the highest sorbic acid resistance on MEA (Fig 1), were also among the strains (DTO006G1, DTO006G7, DTO012A1, DTO012A8, DTO013E5 and DTO013F2) that showed the highest MIC_u_ (Fig 2). DTO013E5 and DTO013F2 were the most resistant even showing growth at the highest tested undissociated sorbate concentration of 21.2 mM, indicating a MIC_u_ > 21.2 mM. The other strains showed a distinctly lower MIC_u_, ranging between 4.2 mM and 9.95 mM. Only strain DTO012A9 showed an intermediate resistance to sorbic acid with an average MIC_u_ of 13.72 mM undissociated sorbic acid.

**Fig 2.**
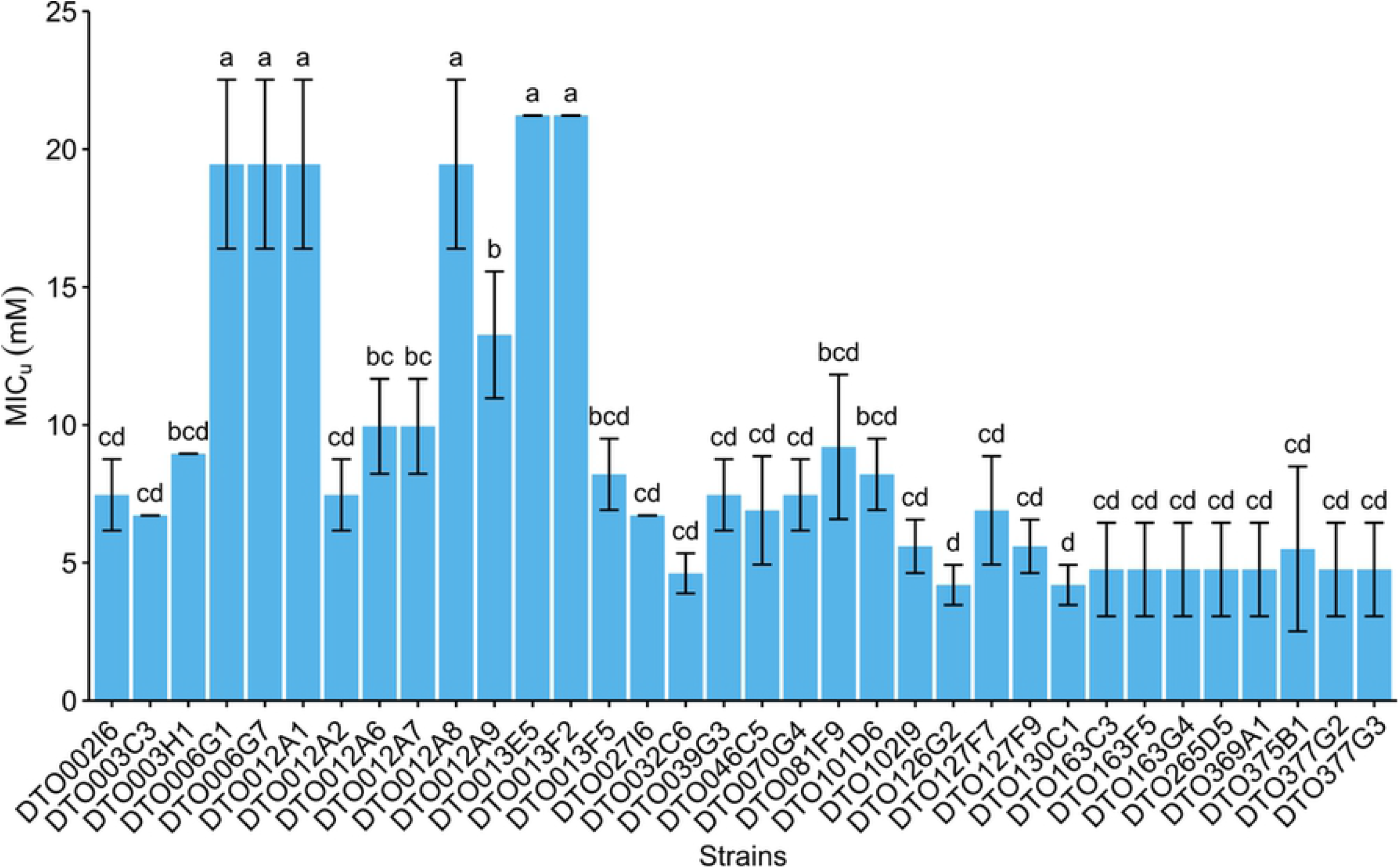
Sorbic acid resistance screening reveals 6 resistant *P. roqueforti* strains. Average undissociated MIC_u_ values of sorbic acid of 34 *P. roqueforti* strains (mM ± standard deviation). Each bar graph represents biological triplicates. Error bars indicate standard deviation and letters indicate significant difference in MIC_u_ (p < 0.05).

### Genome statistics and phylogeny

The genomes of the 34 *P. roqueforti* strains were sequenced. Scaffold count varied between 45 and 1358, assembly length between 26.53 and 31.74 Mb and GC content between 46.85 and 48.44 % (Table 1). The number of predicted genes varied between 9633 and 10644, the number of genes with PFAM domains between 73.11 and 75.38 %, and the number of secondary metabolism gene clusters between 32 and 36. All strains had a BUSCO completeness of >99 %, indicating high quality assembly and gene predictions, except for DTO012A8 with a completeness of 94.83 %. This and the high scaffold count of the DTO012A8 assembly (1358 scaffolds) indicates that its genome assembly is not complete. The strain was kept in the downstream analysis as most other metrics did not differ much compared to the other strains (Table 1). Figure 3 presents a phylogenetic tree of the 34 *P. roqueforti* strains based on 6923 single-copy orthologous genes. Three of the six sorbic acid-resistant strains (DTO012A2, DTO013F2, DTO013E5 and DTO012A8) are similar with DTO012A8 being closely related to those strains. The two other resistant strains are also similar and more distantly related.

**Table 1.**
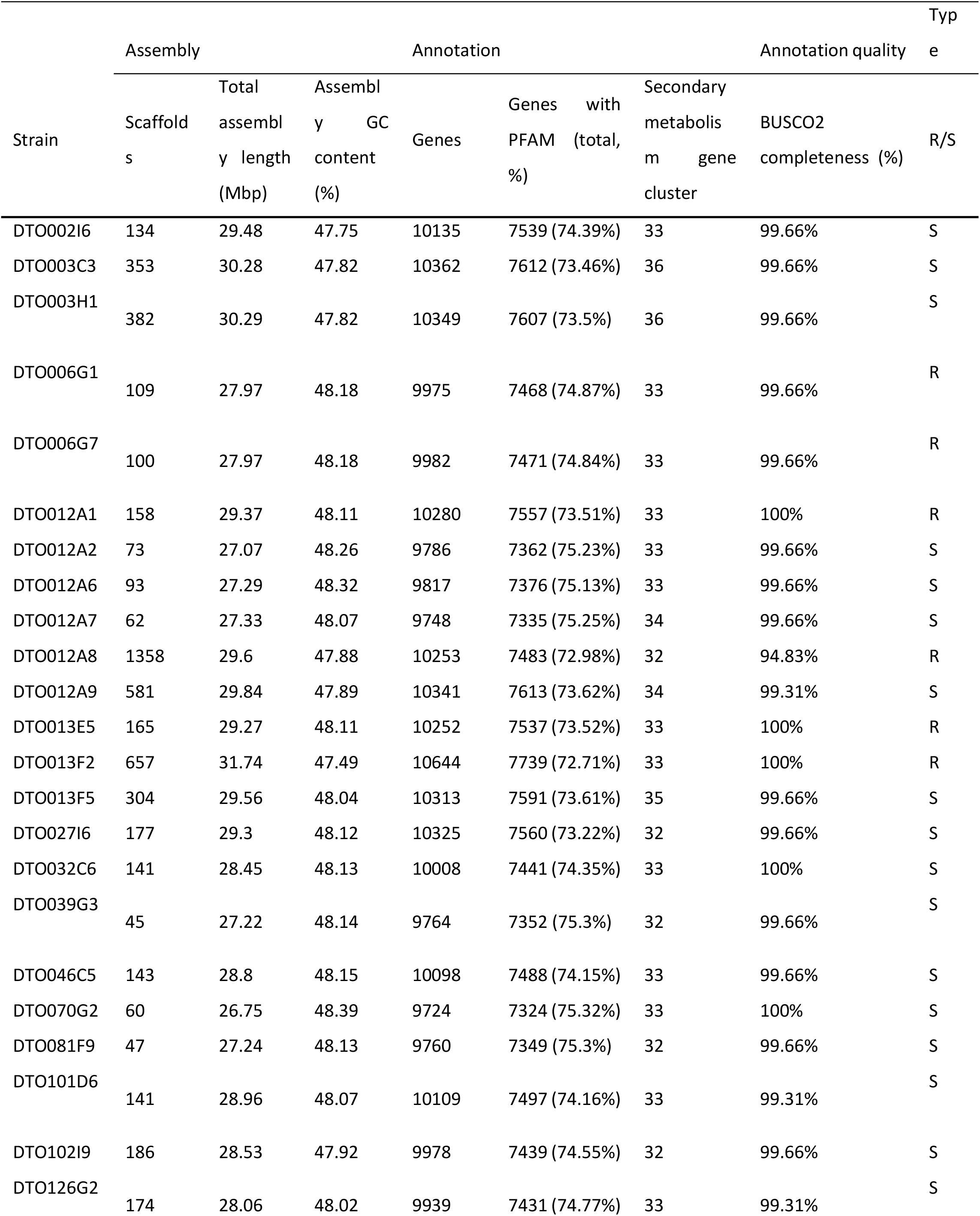
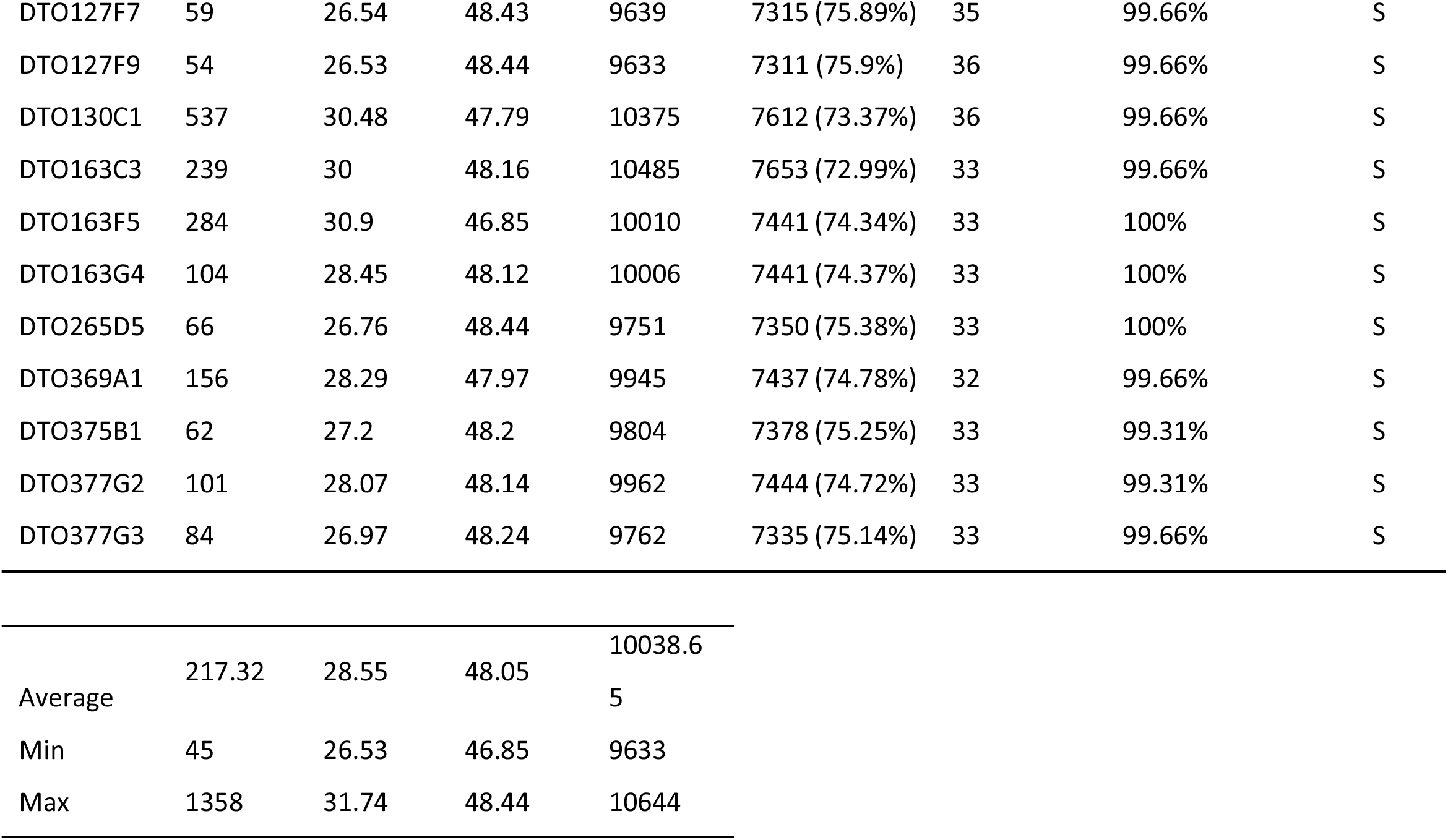
Genome assembly and annotation statistics. The number of scaffolds, assembly length, GC content and genes of the 34 sequenced *P. roqueforti* strains are listed. In addition, the number of genes with a PFAM domain, the number of secondary metabolism gene clusters and the BUSCO completeness are listed. The type column indicates if the strain is sorbic acid-resistant (R) or sorbic acid-sensitive (S).

**Fig 3.**
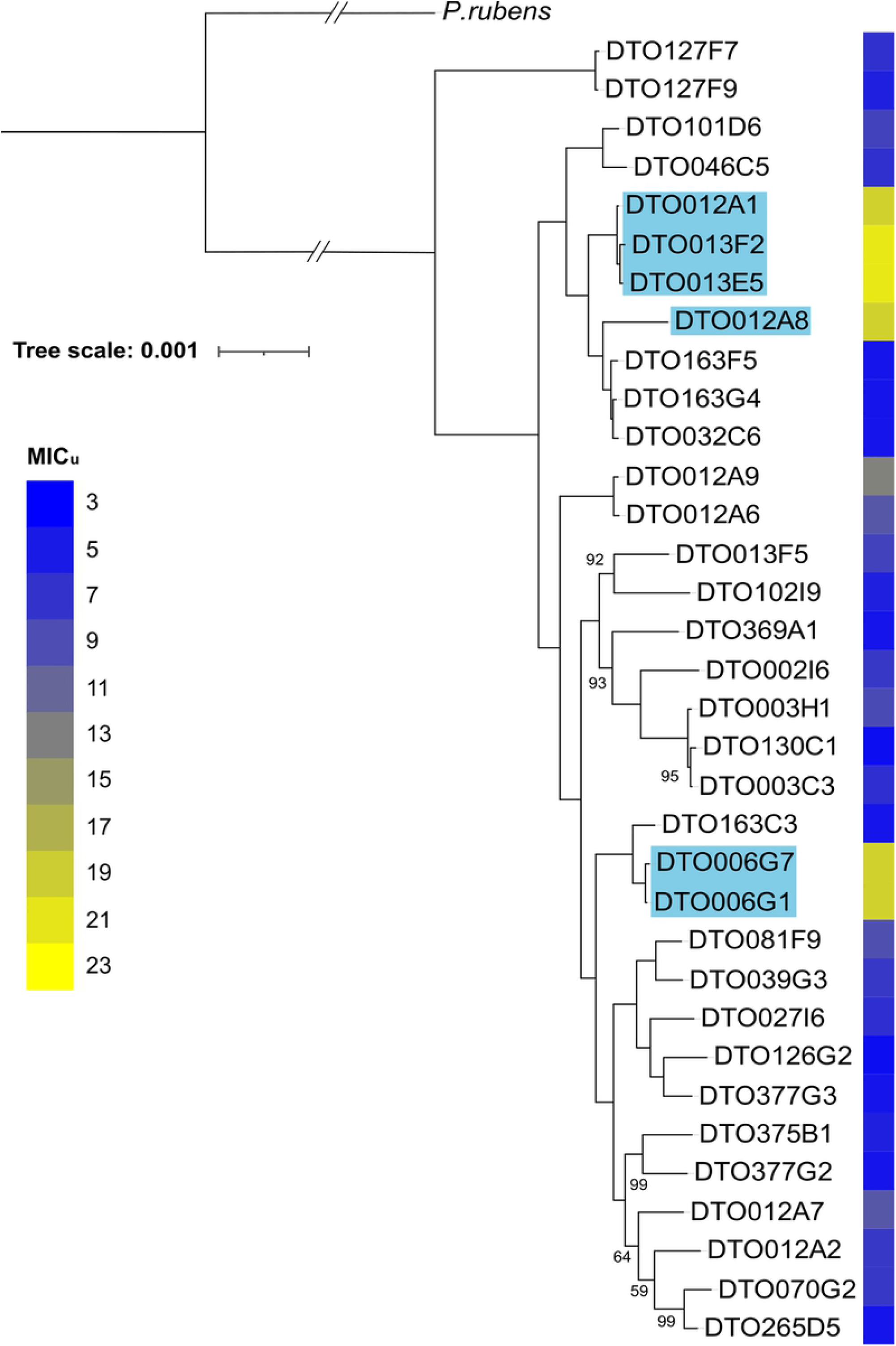
Phylogeny of the *P. roqueforti* strains. Phylogenetic tree of the 34 *P. roqueforti* strains used in this study. Sorbic acid resistance (MIC_u_) is indicated in blue-yellow shading. The tree is based on 6923 single-copy orthologous genes and was constructed using RAxML. *P. rubens* [44] was used as outgroup. Bootstrap values <100 are indicated. Strains containing the SORBUS cluster are highlighted in blue.

### Sorbic acid resistance correlates with a genomic cluster containing genes regulating sorbic acid decarboxylation

The 34 *P. roqueforti* strains were divided into two groups based on their sorbic acid resistance, a group of six resistant (R-type) strains (‘a’, Fig 2) and a group of 28 sensitive (S-type) strains (‘b-d’, Fig 2). Whole genome comparison methods are often based on differences in variants (SNPs), however these methods do no reveal larger missing regions or genes between strains. Hence, a GWAS method was developed to compare whole-genome assemblies based on the MUMmer software. With this method 57 genes unique for the R-type group were identified, of which 51 were present on scaffold 43 of DTO006G7 (Table 2). In addition to the 51 unique genes in this scaffold, it contains 19 genes which are also found completely or in part in some of the S-type strains. The genomic alignment shows the genes on scaffold 43, which is present in the R-type strains (Fig 4). The first 80 kbp of scaffold 43 (protein IDs g12000-g12029) mainly contains hypothetical proteins without predicted function, while the remaining region between 80-180 kbp (g12030 – g12069) contains multiple regions homologous to genes previously reported as related to weak-acid resistance in *A. niger*. Predicted genes orthologous to *padA, cdcA* and *sdrA* of *A. niger* were found alongside each other (g12064-g12066) with respective identities (based on BLAST) of 87 %, 83 % and 53 %. Additional orthologs of *cdcA*, named *cdcB* (g12056) and *cdcC* (g12040) were identified on the same gene cluster as well, with identities of 72 % and 71%, respectively, when compared to *cdcA* of *A. niger*. Another locus (g2591) outside of this cluster also contains a protein homologous to *cdcA* (82 % identity) that is also present in the S-type strains. Similarly, two *padA* orthologs with high BLAST similarities to *padA* of *A. niger* (63 % and 58 %) were identified on the R-type specific cluster and named *padB* (g12032) and *padC* (g12057). In contrast, no homologs of *A. niger sdrA* and *padA* were found outside of the cluster, while 1 and 3 homologs were present inside cluster, respectively. In addition, a transcription factor (g2820 in DTO006G7) orthologous to *warA* was identified outside of the cluster in the genomes of all strains. These results indicate that the R-type strains contain a gene cluster similar to the sorbic acid resistance gene cluster described in *A. niger*, but considerably expanded [9,10]. For further reference, we name this cluster (i.e., scaffold 43 of strain DTO006G7) SORBUS after the tree *Sorbus aucuparia* as sorbic acid has been first isolated from its berries by August Hoffman [19,20]. While SORBUS as a whole is only present in the R-type strains (DTO006G1, DTO006G7, DTO012A2, DTO012A8, DTO013F2, DTO013E5), some S-type strains share up to 5 kbp parts of the sequence, especially in the first 80 kbp of the cluster (Fig 4). It should be noted that the R-type strains were all isolated from non-cheese environments. Based on alignments of the sequencing reads to DTO006G7 we confirmed that out of 35 previously sequenced *P. roqueforti* strains [18], none of the 17 cheese strains contained the SORBUS cluster, while two out of the 18 non-cheese strains contained the SORBUS cluster (Table S1).

**Table 2.**
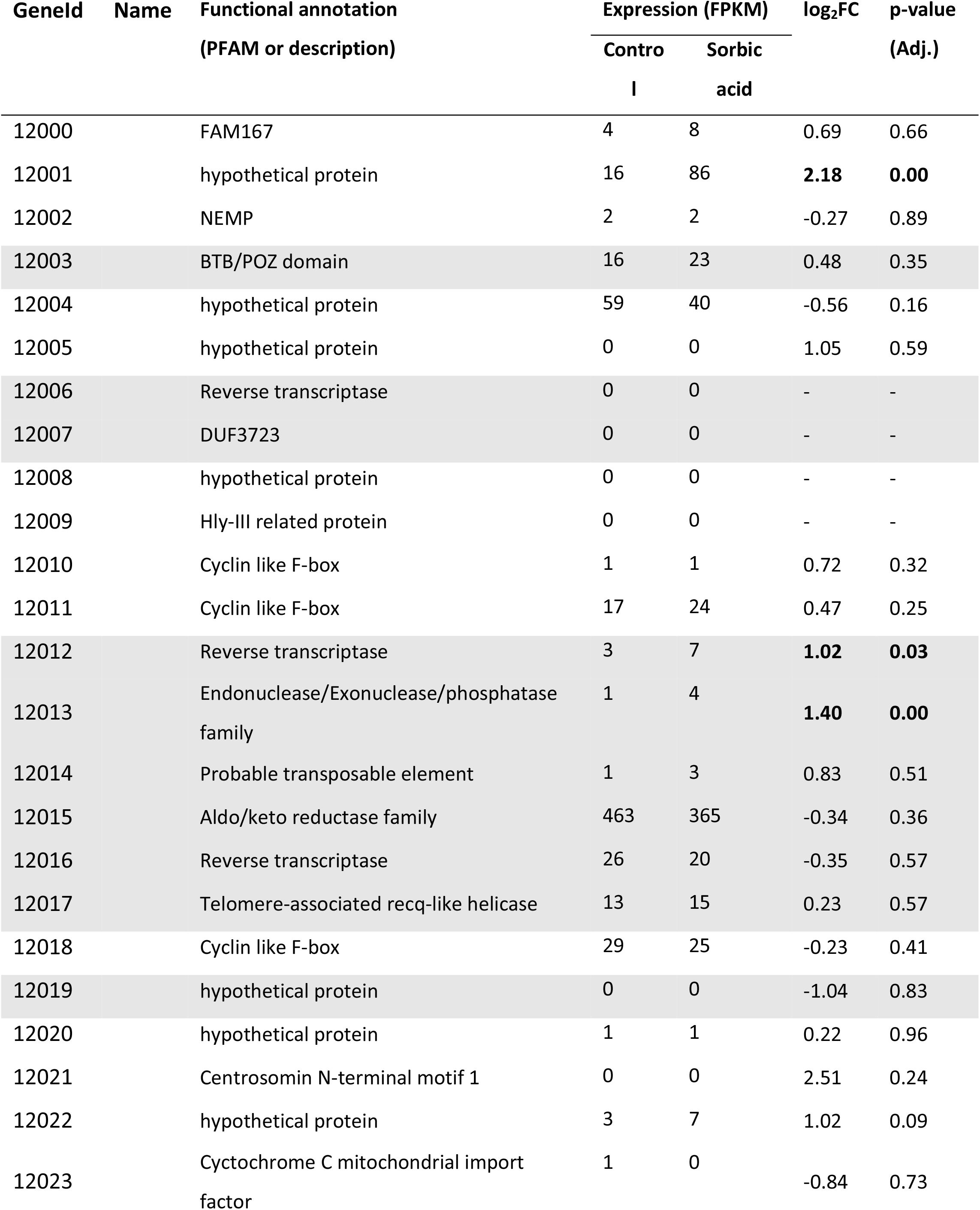
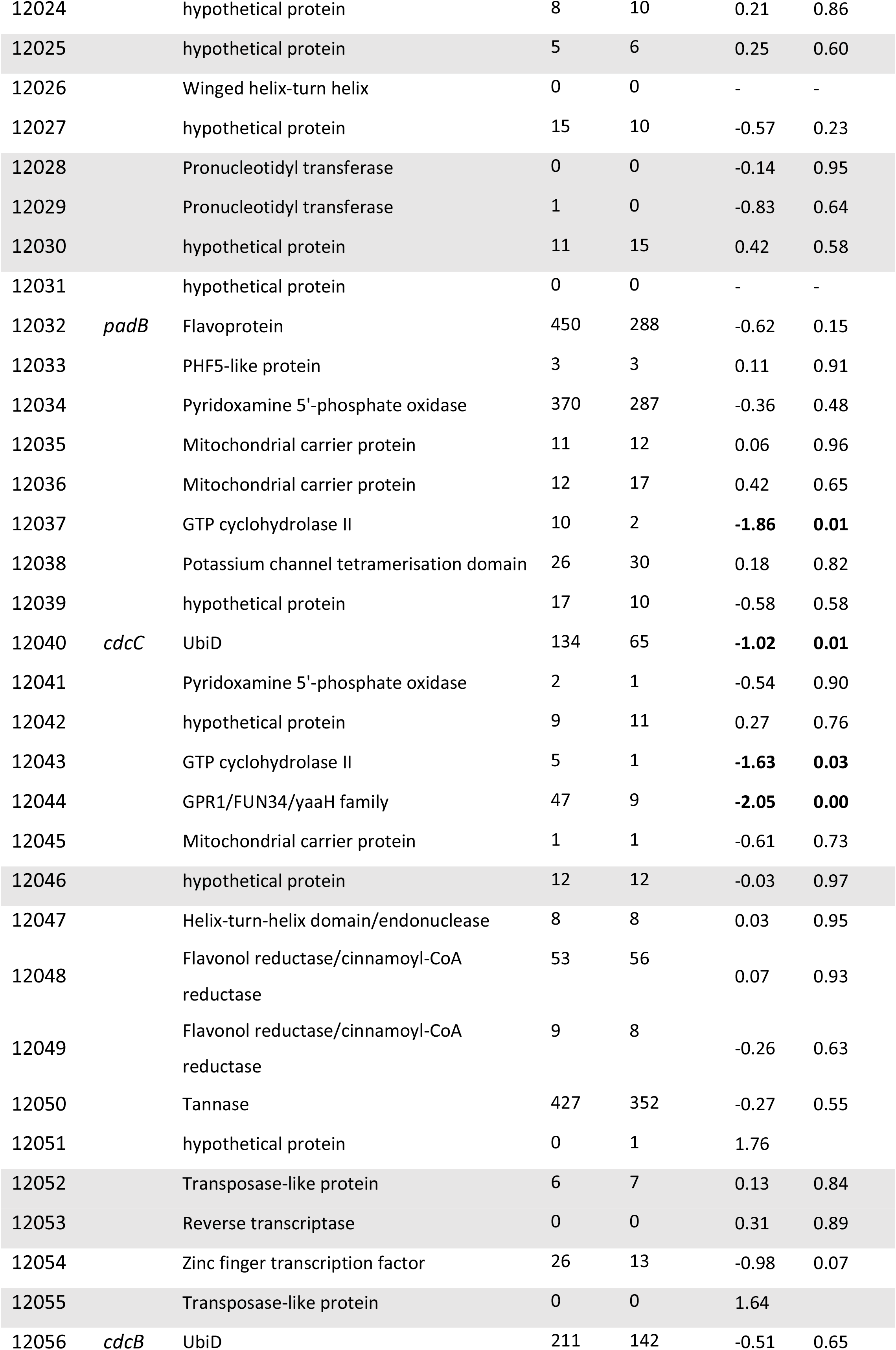
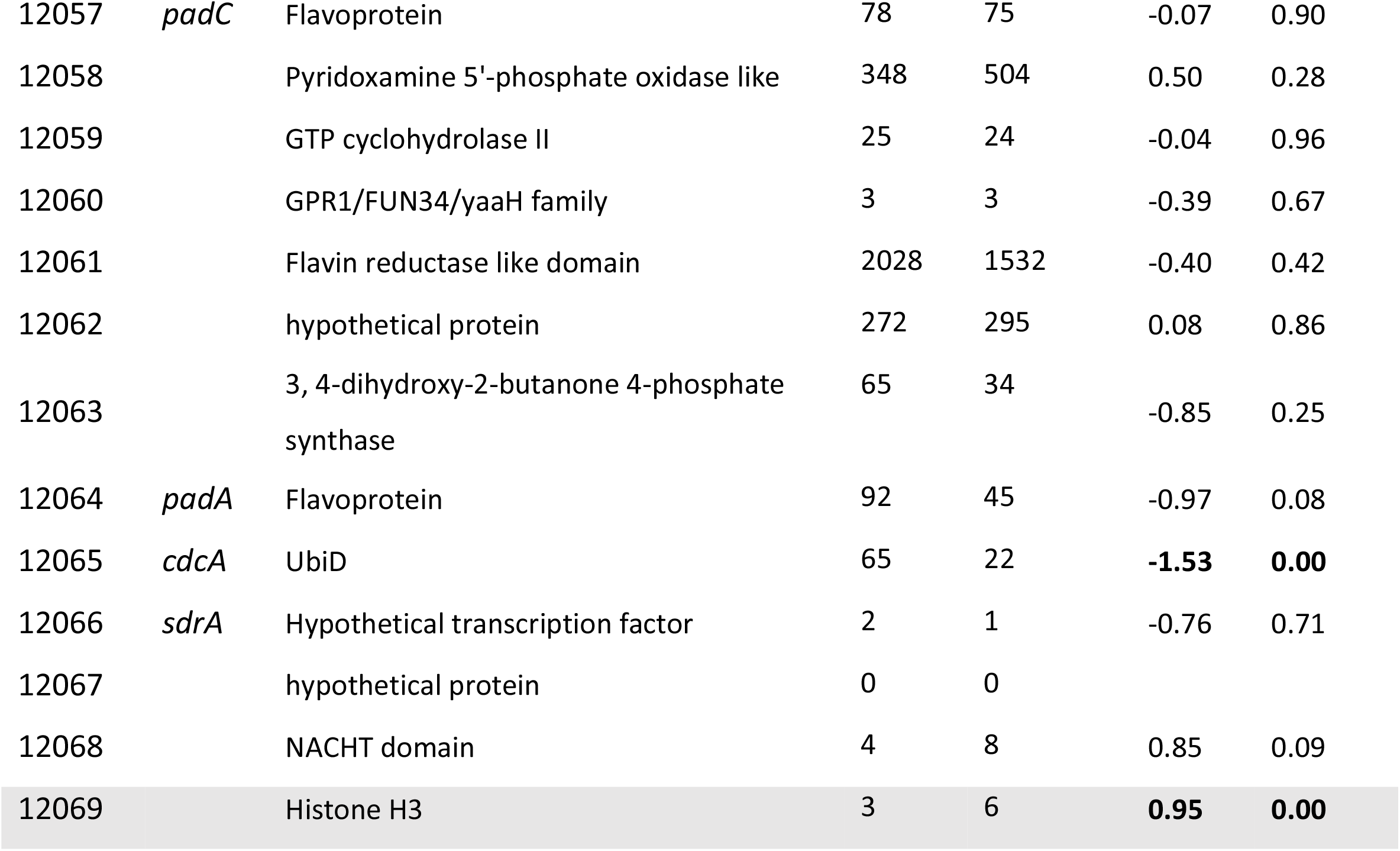
Genes (DTO006G7) located on the SORBUS cluster. Fold change (log_2_FC) of the sorbic acid samples compared to the control is given. Numbers in bold are significantly differently expressed (adjusted p-value < 0.05) and the mean expression (FPKM) of three biological replicates is given per condition (control and sorbic acid). Rows highlighted in grey indicate genes that are not unique for the R-type strains.

| Genelid | Name | Functional annotation<br>(PFAM or description) | Expression (FPKM) | | $\log_2FC$ | p-value<br>(Adj.) |
| --- | --- | --- | --- | --- | --- | --- |
|  |  |  | Contro<br>I | Sorbic<br>acid |  |  |
| 12000 |  | FAM167 | 4 | 8 | 0.69 | 0.66 |
| 12001 |  | hypothetical protein | 16 | 86 | **2.18** | **0.00** |
| 12002 |  | NEMP | 2 | 2 | -0.27 | 0.89 |
| 12003 |  | BTB/POZ domain | 16 | 23 | 0.48 | 0.35 |
| 12004 |  | hypothetical protein | 59 | 40 | -0.56 | 0.16 |
| 12005 |  | hypothetical protein | 0 | 0 | 1.05 | 0.59 |
| 12006 |  | Reverse transcriptase | 0 | 0 | - | - |
| 12007 |  | DUF3723 | 0 | 0 | - | - |
| 12008 |  | hypothetical protein | 0 | 0 | - | - |
| 12009 |  | Hly-III related protein | 0 | 0 | - | - |
| 12010 |  | Cyclin like F-box | 1 | 1 | 0.72 | 0.32 |
| 12011 |  | Cyclin like F-box | 17 | 24 | 0.47 | 0.25 |
| 12012 |  | Reverse transcriptase | 3 | 7 | **1.02** | **0.03** |
| 12013 |  | Endonuclease/Exonuclease/phosphatase family | 1 | 4 | **1.40** | **0.00** |
| 12014 |  | Probable transposable element | 1 | 3 | 0.83 | 0.51 |
| 12015 |  | Aldo/keto reductase family | 463 | 365 | -0.34 | 0.36 |
| 12016 |  | Reverse transcriptase | 26 | 20 | -0.35 | 0.57 |
| 12017 |  | Telomere-associated recq-like helicase | 13 | 15 | 0.23 | 0.57 |
| 12018 |  | Cyclin like F-box | 29 | 25 | -0.23 | 0.41 |
| 12019 |  | hypothetical protein | 0 | 0 | -1.04 | 0.83 |
| 12020 |  | hypothetical protein | 1 | 1 | 0.22 | 0.96 |
| 12021 |  | Centrosomin N-terminal motif 1 | 0 | 0 | 2.51 | 0.24 |
| 12022 |  | hypothetical protein | 3 | 7 | 1.02 | 0.09 |
| 12023 |  | Cytochrome C mitochondrial import factor | 1 | 0 | -0.84 | 0.73 |
| 12024 |  | hypothetical protein | 8 | 10 | 0.21 | 0.86 |
| 12025 |  | hypothetical protein | 5 | 6 | 0.25 | 0.60 |
| 12026 |  | Winged helix-turn helix | 0 | 0 | - | - |
| 12027 |  | hypothetical protein | 15 | 10 | -0.57 | 0.23 |
| 12028 |  | Pronucleotidyl transferase | 0 | 0 | -0.14 | 0.95 |
| 12029 |  | Pronucleotidyl transferase | 1 | 0 | -0.83 | 0.64 |
| 12030 |  | hypothetical protein | 11 | 15 | 0.42 | 0.58 |
| 12031 |  | hypothetical protein | 0 | 0 | - | - |
| 12032 | *padB* | Flavoprotein | 450 | 288 | -0.62 | 0.15 |
| 12033 |  | PHF5-like protein | 3 | 3 | 0.11 | 0.91 |
| 12034 |  | Pyridoxamine 5'-phosphate oxidase | 370 | 287 | -0.36 | 0.48 |
| 12035 |  | Mitochondrial carrier protein | 11 | 12 | 0.06 | 0.96 |
| 12036 |  | Mitochondrial carrier protein | 12 | 17 | 0.42 | 0.65 |
| 12037 |  | GTP cyclohydrolase II | 10 | 2 | **-1.86** | **0.01** |
| 12038 |  | Potassium channel tetramerisation domain | 26 | 30 | 0.18 | 0.82 |
| 12039 |  | hypothetical protein | 17 | 10 | -0.58 | 0.58 |
| 12040 | *cdcC* | UbiD | 134 | 65 | **-1.02** | **0.01** |
| 12041 |  | Pyridoxamine 5'-phosphate oxidase | 2 | 1 | -0.54 | 0.90 |
| 12042 |  | hypothetical protein | 9 | 11 | 0.27 | 0.76 |
| 12043 |  | GTP cyclohydrolase II | 5 | 1 | **-1.63** | **0.03** |
| 12044 |  | GPR1/FUN34/yaaH family | 47 | 9 | **-2.05** | **0.00** |
| 12045 |  | Mitochondrial carrier protein | 1 | 1 | -0.61 | 0.73 |
| 12046 |  | hypothetical protein | 12 | 12 | -0.03 | 0.97 |
| 12047 |  | Helix-turn-helix domain/endonuclease | 8 | 8 | 0.03 | 0.95 |
| 12048 |  | Flavonol reductase/cinnamoyl-CoA reductase | 53 | 56 | 0.07 | 0.93 |
| 12049 |  | Flavonol reductase/cinnamoyl-CoA reductase | 9 | 8 | -0.26 | 0.63 |
| 12050 |  | Tannase | 427 | 352 | -0.27 | 0.55 |
| 12051 |  | hypothetical protein | 0 | 1 | 1.76 |  |
| 12052 |  | Transposase-like protein | 6 | 7 | 0.13 | 0.84 |
| 12053 |  | Reverse transcriptase | 0 | 0 | 0.31 | 0.89 |
| 12054 |  | Zinc finger transcription factor | 26 | 13 | -0.98 | 0.07 |
| 12055 |  | Transposase-like protein | 0 | 0 | 1.64 |  |
| 12056 | *cdcB* | UbiD | 211 | 142 | -0.51 | 0.65 |
| 12057 | *padC* | Flavoprotein | 78 | 75 | -0.07 | 0.90 |
| 12058 |  | Pyridoxamine 5'-phosphate oxidase like | 348 | 504 | 0.50 | 0.28 |
| 12059 |  | GTP cyclohydrolase II | 25 | 24 | -0.04 | 0.96 |
| 12060 |  | GPR1/FUN34/yaaH family | 3 | 3 | -0.39 | 0.67 |
| 12061 |  | Flavin reductase like domain | 2028 | 1532 | -0.40 | 0.42 |
| 12062 |  | hypothetical protein | 272 | 295 | 0.08 | 0.86 |
| 12063 |  | 3, 4-dihydroxy-2-butanone 4-phosphate synthase | 65 | 34 | -0.85 | 0.25 |
| 12064 | *padA* | Flavoprotein | 92 | 45 | -0.97 | 0.08 |
| 12065 | *cdcA* | UbiD | 65 | 22 | **-1.53** | **0.00** |
| 12066 | *sdrA* | Hypothetical transcription factor | 2 | 1 | -0.76 | 0.71 |
| 12067 |  | hypothetical protein | 0 | 0 |  |  |
| 12068 |  | NACHT domain | 4 | 8 | 0.85 | 0.09 |
| 12069 |  | Histone H3 | 3 | 6 | **0.95** | **0.00** |

**Fig 4.**
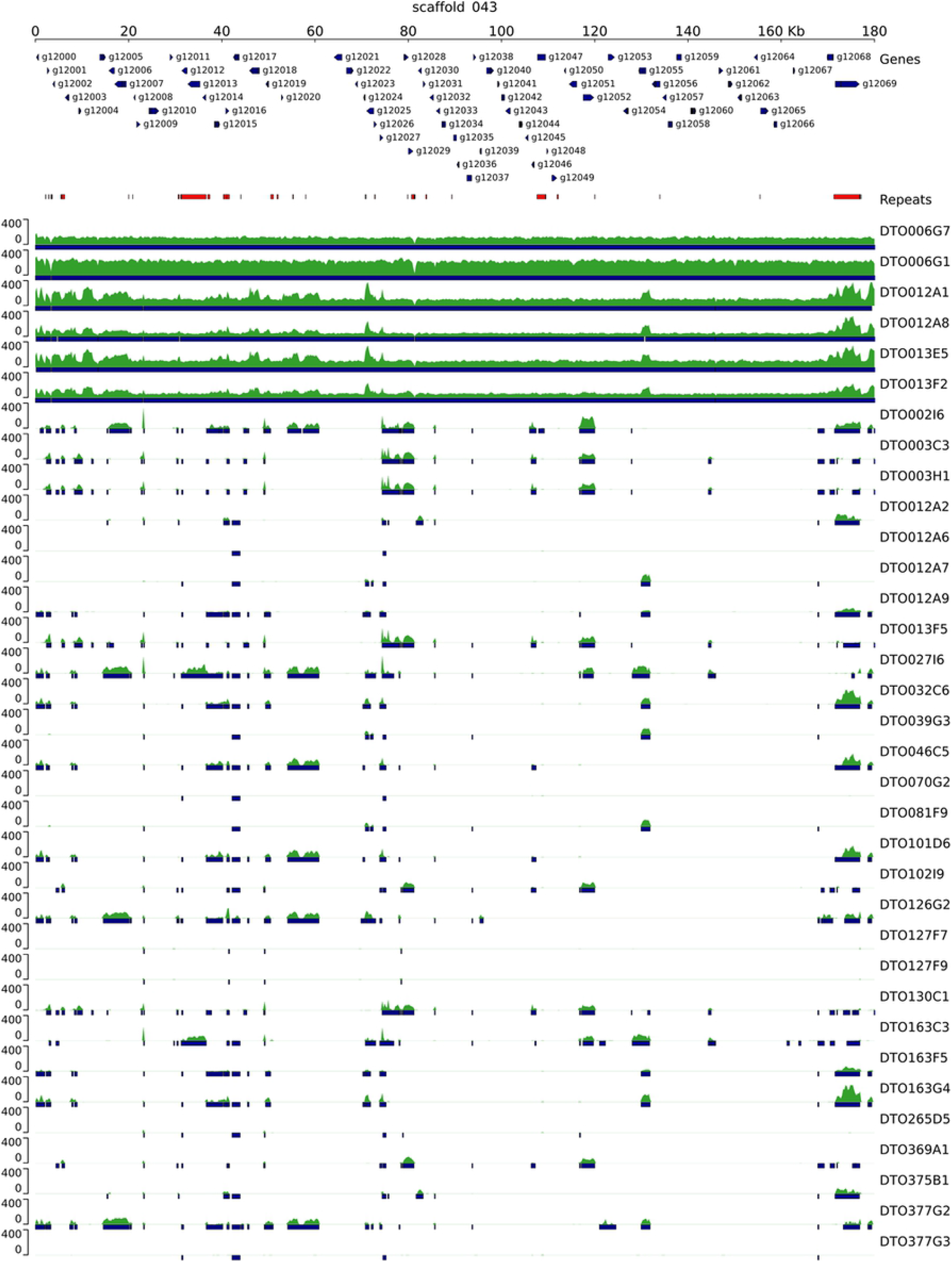
Genome comparison reveals unique gene cluster in R-type strains. Genomic alignment of 33 *P. roqueforti* strains to the SORBUS cluster (scaffold 43 of DTO006G7). Predicted genes are indicated with arrows, repetitive DNA is indicated in red, the sequence read coverage is indicated in the green tracks, and blue bars indicate (partial) overlap with DTO006G7. The complete SORBUS cluster is only present in the R-type strains (DTO006G1, DTO006G7, DTO012A2, DTO012A8, DTO013F2 and DTO013E5).

PLINK [21] was used to identify which SNPs correlate to sorbic acid resistance. This method allowed for the quantitative use of log_10_(MIC_u_) values as input for analysis, as opposed to the approach described above. The SNPs are visualized in a Manhattan plot (Fig 5). SNPs located on genes with a −log_10_(P) > 5 and either a high or moderate impact (SNPeff) were selected (Table 3). This resulted in 338 SNPs in 41 genes. Out of these SNPs, 29 had a ‘high’ impact according to SNPeff and were located in 17 genes. Only six out of these 17 genes with high impact variants (g7017, g8100, g8106, g9942, g9943, g9976) were not located in the SORBUS cluster, the other 11 genes were either among the non-unique genes present on SORBUS, or genes of which less than 90 % of the sequence was found in S-type strains. With a BLAST analysis (protein-protein) and PFAM annotations the functions of the predicted proteins were assessed. This revealed that gene g8100 contains an ankyrin-repeat containing domain and gene g9943 is homologous to a zinc finger C3H1-type domain-containing protein, the putative function of the other three genes could not be assessed (hypothetical proteins). In all cases, the five genes showed high homology (> 99 %) to other *Penicillium* species. The PLINK analysis also revealed SNPs in two genes encoding proteins with a putative transmembrane transporter (g216) or cation transporter (g296) function.

**Table 3.** Genes (DTO006G7) containing SNPs associated with sorbic acid resistance. Only SNPs with a −log_10_(p-value) > 5 and a moderate or high impact as determined by SNPeff are listed. Grey shading indicates overlap with sequence repeats.

| GeneID | SNP impact |  | Effect | PFAM annotation |
| --- | --- | --- | --- | --- |
|  | Moderate | High |  |  |
| g103 | 1 | 0 |  | hypothetical protein |
| g216 | 1 | 0 |  | ABC transporter transmembrane region |
| g235 | 1 | 0 |  | STAG domain |
| g296 | 5 | 0 |  | Cation transporter/ATPase, N-terminus |
| g312 | 1 | 0 |  | Probable molybdopterin binding domain |
| g313 | 1 | 0 |  | DDHD domain |
| g314 | 1 | 0 |  | hypothetical protein |
| g315 | 1 | 0 |  | FAD binding domain |
| g7015 | 1 | 1 | Stop lost & splice region variant | hypothetical protein |
| g8100 | 0 | 1 | Stop gained | Ankyrin repeats (3 copies) |
| g8101 | 3 | 0 |  | DDE superfamily endonuclease |
| g8104 | 7 | 0 |  | hypothetical protein |
| g8105 | 10 | 0 |  | hypothetical protein |
| g8106 | 3 | 1 | Stop gained | hypothetical protein |
| g8107 | 26 | 0 |  | Ankyrin repeats (3 copies) |
| g12002 | 2 | 0 |  | NEMP |
| g12005 | 9 | 1 | Frameshift variant | hypothetical protein |
| g12006 | 42 | 2 | Frameshift variant & Stop gained | Reverse transcriptase |
| g12007 | 8 | 0 |  | Protein of unknown function (DUF3723) |
| g12013 | 48 | 3 | Frameshift variants | Endonuclease/Exonuclease/phosphatase family |
| g12014 | 8 | 1 | Stop lost & splice region variant | Probable transposable element |
| g12016 | 2 | 0 |  | Reverse transcriptase |
| g12019 | 11 | 0 |  | hypothetical protein |
| g12025 | 9 | 1 | Stop gained | hypothetical protein |
| g12028 | 18 | 1 | Stop lost & splice region variant | Pronucleotidyl transferase |
| g12029 | 26 | 6 | Stop gained, Stop lost & splice variant | Pronucleotidyl transferase |
| g12030 | 16 | 2 | Frameshift variant & Stop lost | hypothetical protein |
| g12046 | 1 | 0 |  | hypothetical protein |
| g12052 | 4 | 1 | Stop gained | Transposase-like protein |
| g12053 | 1 | 0 |  | Reverse transcriptase |
| g12054 | 4 | 0 |  | hypothetical protein |
| g12055 | 6 | 2 | Frameshift variant & Stop lost | Transposase-like protein |
| g12069 | 7 | 1 | Frameshift variant | Histone H3 |
| g9749 | 7 | 0 |  | hypothetical protein |
| g9942 | 1 | 2 | Stop gained | hypothetical protein |
| g9943 | 2 | 2 | Stop gained | AAA domain |
| g9944 | 8 | 0 |  | hypothetical protein |
| g9945 | 4 | 0 |  | ATPase family associated with various cellular activities (AAA) |
| g9961 | 1 | 0 |  | hypothetical protein |
| g9966 | 2 | 0 |  | Endonuclease-reverse transcriptase |
| g9976 | 0 | 1 | Start lost | hypothetical protein |
| **TOTAL** | **309** | **29** |  |  |

**Fig 5.**
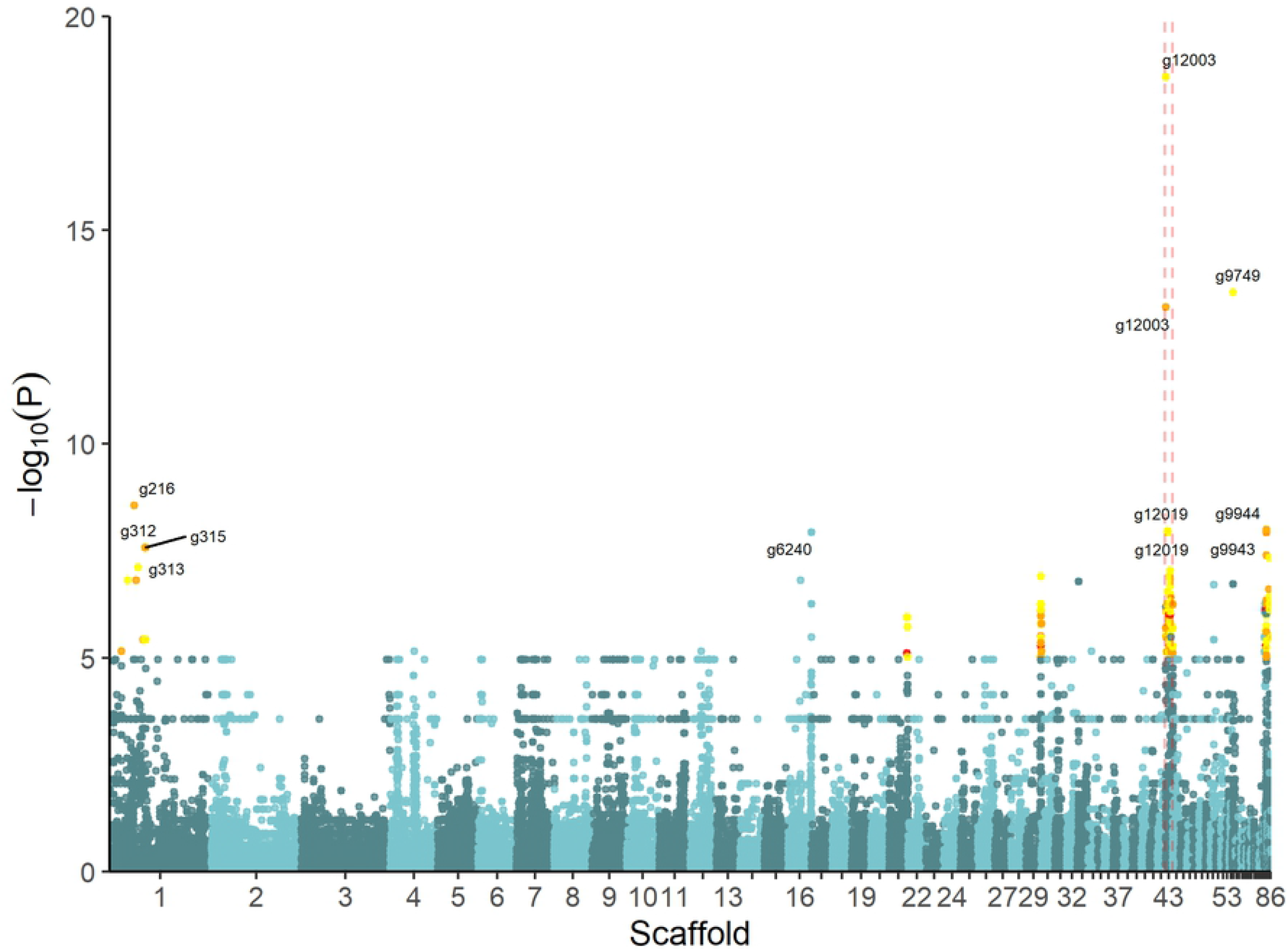
Manhattan plot shows SNPs associated with sorbic acid resistance. Scaffolds are listed on the x-axis, while the y-axis display the significance of the association (−log_10_(p-value)). Yellow, orange and red dots indicate ‘low’, ‘moderate’ or ‘high’ impact SNPs as determined by SNPeff, respectively. The GeneIDs associated with the SNPs with a −log_10_(p-value) > 7.5 are indicated. The SORBUS cluster is located between the dashed lines.

To investigate the evolutionary origin of this cluster, presence of five PFAM domains (from g12060, g12061 and g12063-g12065) that are present on the SORBUS cluster was analysed in 32 Aspergilli and Penicillia as well as the 34 *P. roqueforti* strains (Table 4). These PFAM domains encode a putative GPR1/FUN34/yaaH family (g12060), a flavin reductase like domain (g12061), 3, 4-dihydroxy-2-butanone 4-phosphate synthase (g12063), a flavoprotein (g12064, *padA*) and a UbiD domain (g12065, *cdcA*). These domains are selected as they are clustered and because of their predicted role. The first domain has been associated with acetic acid sensitivity in *S. cerevisiae* [22], while the latter four domains are part of the gene cluster described in *A. niger* [9]. Genes containing these domains were aligned using MAFFT and alignments were used to construct a phylogenetic tree per domain. Based on these phylogenetic trees the relatedness of the five genes of the SORBUS cluster was assessed. This revealed that the SORBUS genes g12061, g12064 and g12065 cluster more closely with several *Aspergillus* species (Table 4), whereas the PFAM domains from g12060 and g12063 did not cluster with any of the species included. In addition, the PFAMs present in the core genome (present both in S- and R-type strains) aligned more closely to *P. digitatum* or *P. oxalicum*. Furthermore, two genes with homology to transposase-like proteins (g12052 and g12055) and several reverse-transcriptase domains were identified in the SORBUS cluster.

**Table 4.**
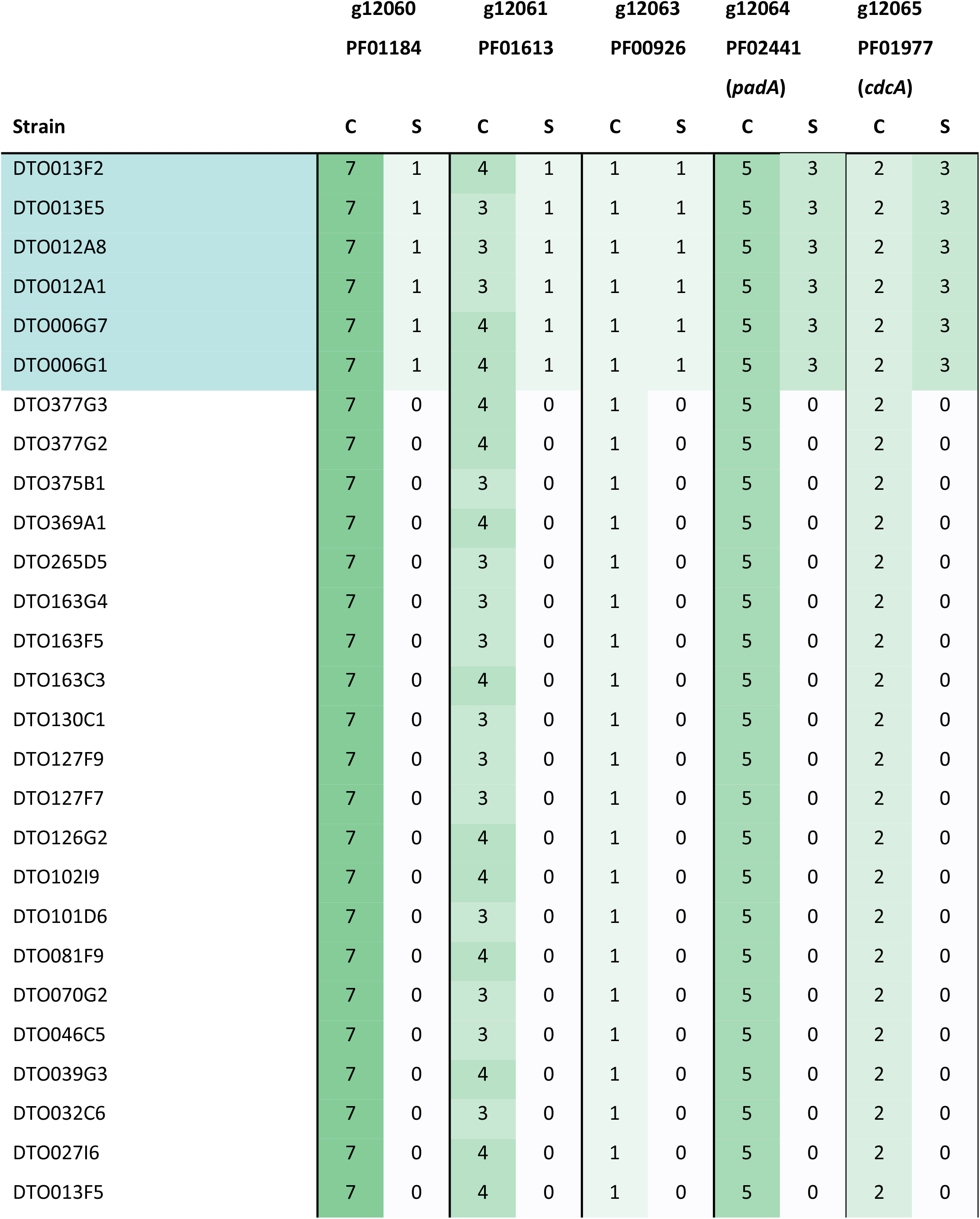
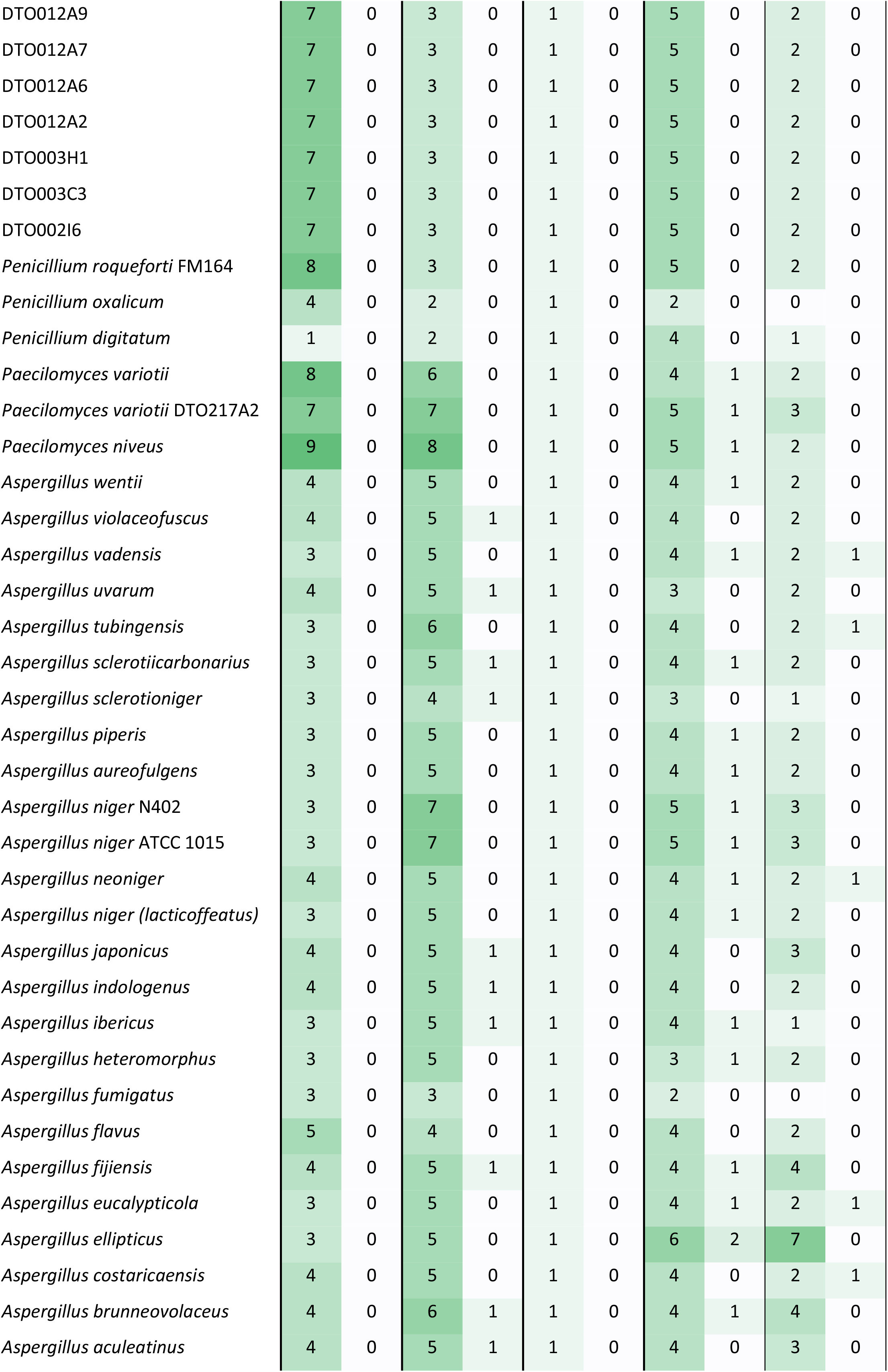
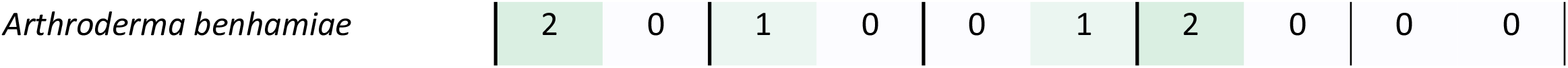
Number of genes containing PFAM domains corresponding to g12060, g12061 and g12063-g12065 (PF01184, PF01613, PF00926, PF02441, PF01977) based on phylogenetic trees constructed with their respective PFAM domains. Top six strains are R-type *P. roqueforti* strains containing the SORBUS cluster. C (CORE) indicates if the domains aligned closely to the PFAMs not unique for the SORBUS cluster or did not align to *P. roqueforti*, S (SORBUS) indicates the number of PFAM domains which aligned closely to PFAMs originated from SORBUS.

| Strain | g12060 |  | g12061 |  | g12063 |  | g12064 |  | g12065 |  |
| --- | --- | --- | --- | --- | --- | --- | --- | --- | --- | --- |
|  | PF01184 |  | PF01613 |  | PF00926 |  | PF02441 |  | PF01977 |  |
|  |  |  |  |  |  |  | (padA) |  | (cdcA) |  |
|  | C | S | C | S | C | S | C | S | C | S |
| DT0013F2 | 7 | 1 | 4 | 1 | 1 | 1 | 5 | 3 | 2 | 3 |
| DT0013E5 | 7 | 1 | 3 | 1 | 1 | 1 | 5 | 3 | 2 | 3 |
| DT0012A8 | 7 | 1 | 3 | 1 | 1 | 1 | 5 | 3 | 2 | 3 |
| DT0012A1 | 7 | 1 | 3 | 1 | 1 | 1 | 5 | 3 | 2 | 3 |
| DT0006G7 | 7 | 1 | 4 | 1 | 1 | 1 | 5 | 3 | 2 | 3 |
| DT0006G1 | 7 | 1 | 4 | 1 | 1 | 1 | 5 | 3 | 2 | 3 |
| DT0377G3 | 7 | 0 | 4 | 0 | 1 | 0 | 5 | 0 | 2 | 0 |
| DT0377G2 | 7 | 0 | 4 | 0 | 1 | 0 | 5 | 0 | 2 | 0 |
| DT0375B1 | 7 | 0 | 3 | 0 | 1 | 0 | 5 | 0 | 2 | 0 |
| DT0369A1 | 7 | 0 | 4 | 0 | 1 | 0 | 5 | 0 | 2 | 0 |
| DT0265D5 | 7 | 0 | 3 | 0 | 1 | 0 | 5 | 0 | 2 | 0 |
| DT0163G4 | 7 | 0 | 3 | 0 | 1 | 0 | 5 | 0 | 2 | 0 |
| DT0163F5 | 7 | 0 | 3 | 0 | 1 | 0 | 5 | 0 | 2 | 0 |
| DT0163C3 | 7 | 0 | 4 | 0 | 1 | 0 | 5 | 0 | 2 | 0 |
| DT0130C1 | 7 | 0 | 3 | 0 | 1 | 0 | 5 | 0 | 2 | 0 |
| DT0127F9 | 7 | 0 | 3 | 0 | 1 | 0 | 5 | 0 | 2 | 0 |
| DT0127F7 | 7 | 0 | 3 | 0 | 1 | 0 | 5 | 0 | 2 | 0 |
| DT0126G2 | 7 | 0 | 4 | 0 | 1 | 0 | 5 | 0 | 2 | 0 |
| DT0102I9 | 7 | 0 | 4 | 0 | 1 | 0 | 5 | 0 | 2 | 0 |
| DT0101D6 | 7 | 0 | 3 | 0 | 1 | 0 | 5 | 0 | 2 | 0 |
| DT0081F9 | 7 | 0 | 4 | 0 | 1 | 0 | 5 | 0 | 2 | 0 |
| DT0070G2 | 7 | 0 | 3 | 0 | 1 | 0 | 5 | 0 | 2 | 0 |
| DT0046C5 | 7 | 0 | 3 | 0 | 1 | 0 | 5 | 0 | 2 | 0 |
| DT0039G3 | 7 | 0 | 4 | 0 | 1 | 0 | 5 | 0 | 2 | 0 |
| DT0032C6 | 7 | 0 | 3 | 0 | 1 | 0 | 5 | 0 | 2 | 0 |
| DT0027I6 | 7 | 0 | 4 | 0 | 1 | 0 | 5 | 0 | 2 | 0 |
| DT0013F5 | 7 | 0 | 4 | 0 | 1 | 0 | 5 | 0 | 2 | 0 |
| DTO012A9 | 7 | 0 | 3 | 0 | 1 | 0 | 5 | 0 | 2 | 0 |
| DTO012A7 | 7 | 0 | 3 | 0 | 1 | 0 | 5 | 0 | 2 | 0 |
| DTO012A6 | 7 | 0 | 3 | 0 | 1 | 0 | 5 | 0 | 2 | 0 |
| DTO012A2 | 7 | 0 | 3 | 0 | 1 | 0 | 5 | 0 | 2 | 0 |
| DTO003H1 | 7 | 0 | 3 | 0 | 1 | 0 | 5 | 0 | 2 | 0 |
| DTO003C3 | 7 | 0 | 3 | 0 | 1 | 0 | 5 | 0 | 2 | 0 |
| DTO002I6 | 7 | 0 | 3 | 0 | 1 | 0 | 5 | 0 | 2 | 0 |
| *Penicillium roqueforti* FM164 | 8 | 0 | 3 | 0 | 1 | 0 | 5 | 0 | 2 | 0 |
| *Penicillium oxalicum* | 4 | 0 | 2 | 0 | 1 | 0 | 2 | 0 | 0 | 0 |
| *Penicillium digitatum* | 1 | 0 | 2 | 0 | 1 | 0 | 4 | 0 | 1 | 0 |
| *Paecilomyces variotii* | 8 | 0 | 6 | 0 | 1 | 0 | 4 | 1 | 2 | 0 |
| *Paecilomyces variotii* DTO217A2 | 7 | 0 | 7 | 0 | 1 | 0 | 5 | 1 | 3 | 0 |
| *Paecilomyces niveus* | 9 | 0 | 8 | 0 | 1 | 0 | 5 | 1 | 2 | 0 |
| *Aspergillus wentii* | 4 | 0 | 5 | 0 | 1 | 0 | 4 | 1 | 2 | 0 |
| *Aspergillus violaceofuscus* | 4 | 0 | 5 | 1 | 1 | 0 | 4 | 0 | 2 | 0 |
| *Aspergillus vadensis* | 3 | 0 | 5 | 0 | 1 | 0 | 4 | 1 | 2 | 1 |
| *Aspergillus uvarum* | 4 | 0 | 5 | 1 | 1 | 0 | 3 | 0 | 2 | 0 |
| *Aspergillus tubingensis* | 3 | 0 | 6 | 0 | 1 | 0 | 4 | 0 | 2 | 1 |
| *Aspergillus sclerotiicarbonarius* | 3 | 0 | 5 | 1 | 1 | 0 | 4 | 1 | 2 | 0 |
| *Aspergillus sclerotioniger* | 3 | 0 | 4 | 1 | 1 | 0 | 3 | 0 | 1 | 0 |
| *Aspergillus piperis* | 3 | 0 | 5 | 0 | 1 | 0 | 4 | 1 | 2 | 0 |
| *Aspergillus aureofulgens* | 3 | 0 | 5 | 0 | 1 | 0 | 4 | 1 | 2 | 0 |
| *Aspergillus niger* N402 | 3 | 0 | 7 | 0 | 1 | 0 | 5 | 1 | 3 | 0 |
| *Aspergillus niger* ATCC 1015 | 3 | 0 | 7 | 0 | 1 | 0 | 5 | 1 | 3 | 0 |
| *Aspergillus neoniger* | 4 | 0 | 5 | 0 | 1 | 0 | 4 | 1 | 2 | 1 |
| *Aspergillus niger (lacticoffeatus)* | 3 | 0 | 5 | 0 | 1 | 0 | 4 | 1 | 2 | 0 |
| *Aspergillus japonicus* | 4 | 0 | 5 | 1 | 1 | 0 | 4 | 0 | 3 | 0 |
| *Aspergillus indologenus* | 4 | 0 | 5 | 1 | 1 | 0 | 4 | 0 | 2 | 0 |
| *Aspergillus ibericus* | 3 | 0 | 5 | 1 | 1 | 0 | 4 | 1 | 1 | 0 |
| *Aspergillus heteromorphus* | 3 | 0 | 5 | 0 | 1 | 0 | 3 | 1 | 2 | 0 |
| *Aspergillus fumigatus* | 3 | 0 | 3 | 0 | 1 | 0 | 2 | 0 | 0 | 0 |
| *Aspergillus flavus* | 5 | 0 | 4 | 0 | 1 | 0 | 4 | 0 | 2 | 0 |
| *Aspergillus fijiensis* | 4 | 0 | 5 | 1 | 1 | 0 | 4 | 1 | 4 | 0 |
| *Aspergillus eucalypticola* | 3 | 0 | 5 | 0 | 1 | 0 | 4 | 1 | 2 | 1 |
| *Aspergillus ellipticus* | 3 | 0 | 5 | 0 | 1 | 0 | 6 | 2 | 7 | 0 |
| *Aspergillus costaricaensis* | 4 | 0 | 5 | 0 | 1 | 0 | 4 | 0 | 2 | 1 |
| *Aspergillus brunneovolaceus* | 4 | 0 | 6 | 1 | 1 | 0 | 4 | 1 | 4 | 0 |
| *Aspergillus aculeatinus* | 4 | 0 | 5 | 1 | 1 | 0 | 4 | 0 | 3 | 0 |

*Arthroderma benhamiae*
|  |  |  |  |  |  |  |  |  |  |
|---|---|---|---|---|---|---|---|---|---|
| 2 | 0 | 1 | 0 | 0 | 1 | 2 | 0 | 0 | 0 |

### Genome-wide expression profiles of a sorbic acid-sensitive and sorbic acid-resistant strain

A genome-wide expression profile was performed on a sorbic acid-sensitive strain (DTO377G3) and a sorbic acid-resistant strain (DTO006G7) grown on MEB in the presence or absence of 3 mM sorbic acid. The sequence reads were aligned to the assemblies of these two *P. roqueforti* strains. Gene expression values were calculated and differentially expressed genes were identified (Table S4). The expression profiles of the biological replicates were similar, demonstrated by their clustering in the PCA plot (Fig 6). Combined, PC1 and PC2 explain 90 % of the variation observed. The samples treated with sorbic acid separate from the control samples on Y-axis while the differences between the strains are separated by on the X-axis.

**Fig 6.**
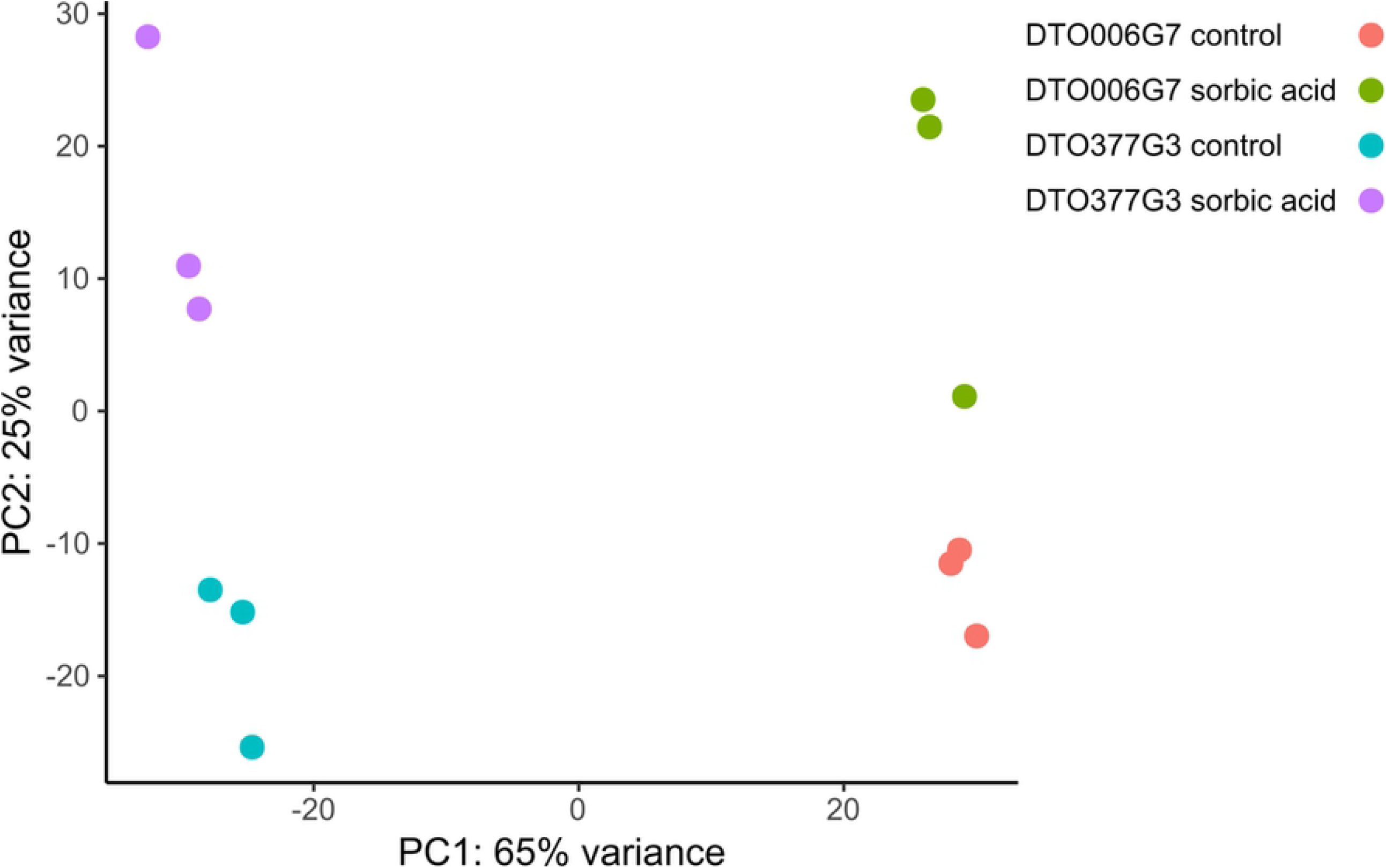
Principle component analysis demonstrates clustering of sorbic acid-exposed strains. Each dot represents a biological replicate. PC1 and PC2 together describe 90% of the variation. The sample grown on sorbic acid separate on the Y-axis and the two strains are separated on the X-axis.

Genes that are either up- or down-regulated in both the R-type and S-type strain when exposed to sorbic acid might be involved in a general response to sorbic acid stress in *P. roqueforti*. Venn diagrams were constructed revealing that 33 genes were significantly up-regulated in both the S-type and R-type strain (Fig 7A). An enrichment analysis revealed that the functional annotation terms ‘secretion signal’ and ‘small secreted protein’ are over-represented in these genes. Among the 21 shared down-regulated genes (Fig 7B) the NmrA-like family and NAD(P)H-binding domains were over-represented. Table S5 lists all genes present in both shared pools. The expression of the SORBUS genes (g12000-g12069) was analysed. This revealed that nine out of the 70 genes were significantly differentially expressed (Table 2), including two out of the three genes homologous to *cdcA* that were lower expressed (log_2_FC = −1.5) in the presence of sorbic acid. On the SORBUS cluster g12061 (with a flavin-reductase like domain) was the highest expressed gene, reaching 2000 FPKM in the control condition.

**Fig 7.**
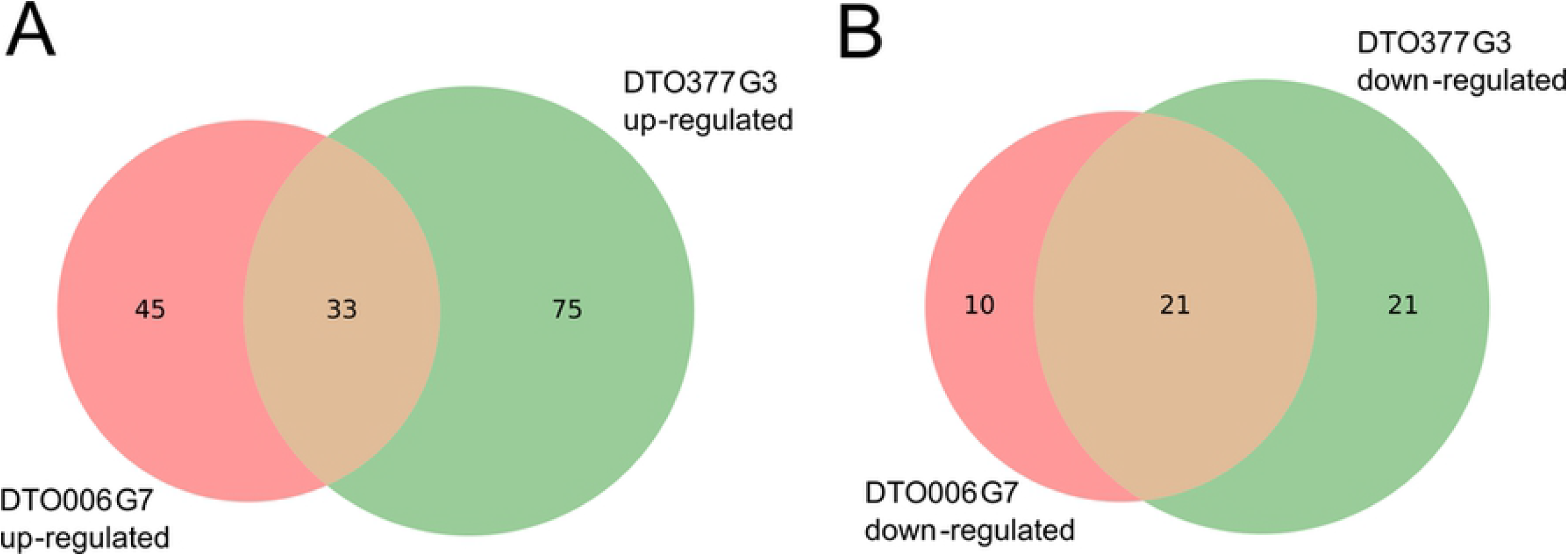
Venn diagram of differentially expressed genes in the presence of sorbic acid. Up- (A) or down-regulated (B) in R-type DTO006G7 and S-type DTO377G3 when exposed to sorbic acid compared to the control in the absence of this weak acid. Genes were considered up- or down-regulated when the log_2_FC > 2, p-value < 0.05 and FPKM >10.

### Partial knock-out confirms role of SORBUS in sorbic acid resistance

Strains *P. roqueforti* DTO013F2Δ*kusA* SC1 and SC2 were obtained lacking part of the SORBUS cluster. Nanopore sequencing was used to investigate how much of the SORBUS cluster was removed in these two transformants (Fig 8). This revealed that the deletions were larger than the 93 kbp region. A 105 and a 131 kbp part of the cluster was removed in DTO013F2 Δ*kusA* SC1 and SC2, respectively. Detailed analysis of the nanopore reads revealed that the smaller fragments within the SORBUS cluster present in SC1 and SC2 strains consisted of highly dissimilar sequences, indicating these are parts of repetitive DNA (Fig 8). Both SC strains had a reduced resistance to sorbic acid when compared to DTO013F2 and DTO013F2 Δ*kusA* with a MIC_u_ similar to the S-type strain DTO377G3 (Fig 9). This shows that part of the SORBUS cluster is involved in sorbic acid stress mitigation.

**Fig 8.**
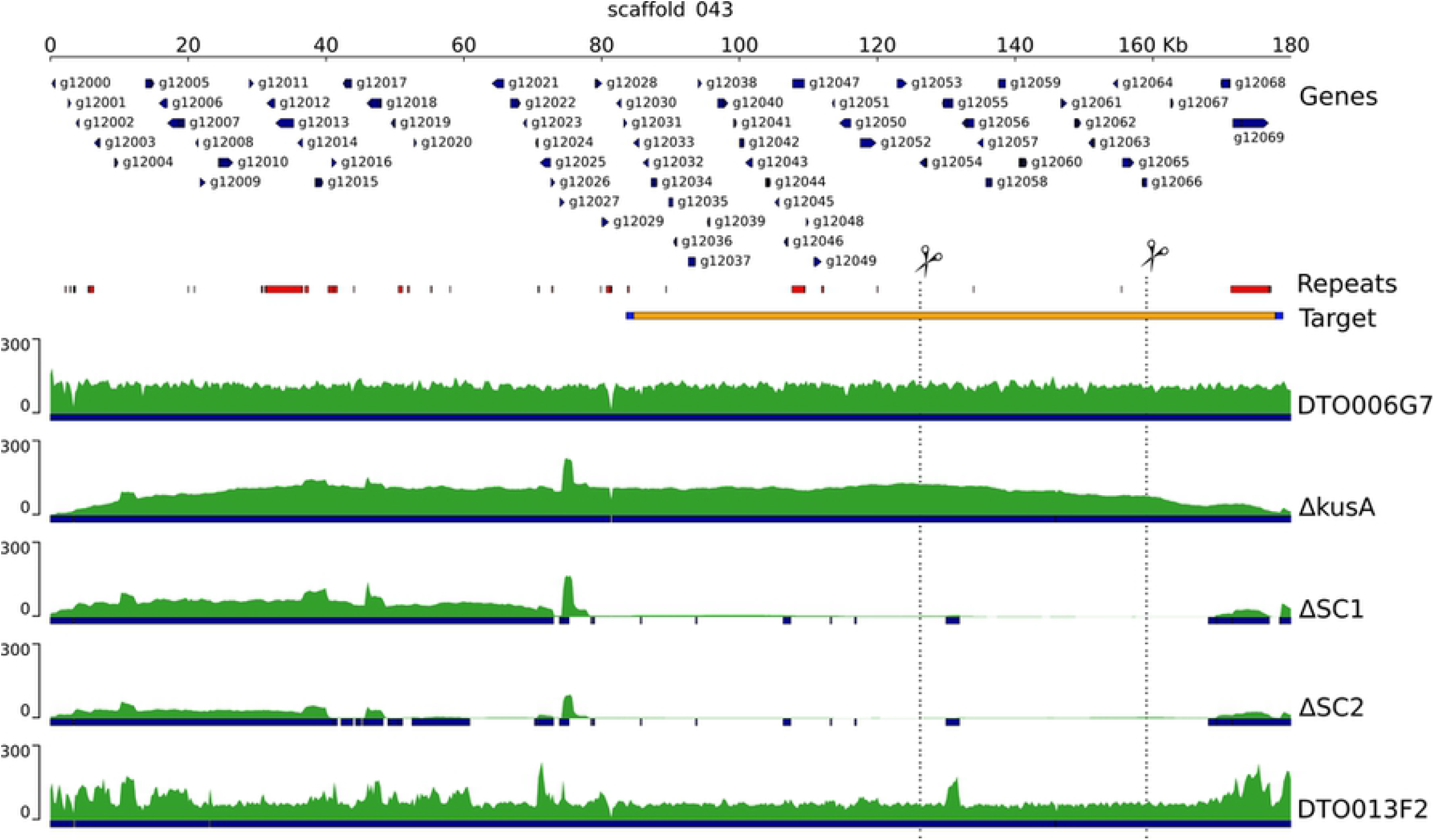
Schematic overview of SORBUS sequences of the knockouts strains. Strains DTO006G7, DTO013F2 and knock-out strains DTO013F2 Δ*kusA*, SC1 and SC2 are depicted. Predicted genes are indicated with arrows, repetitive DNA is indicated in red, the sequence read coverage is indicated in the green tracks, and blue bars indicate (partial) overlap with DTO006G7. The orange bar indicates the targeted knock-out region (target), flanked by the 5’ and 3’ ends (in blue). Dotted lines and scissors indicate the loci targeted by the sgRNAs.

**Fig 9.**
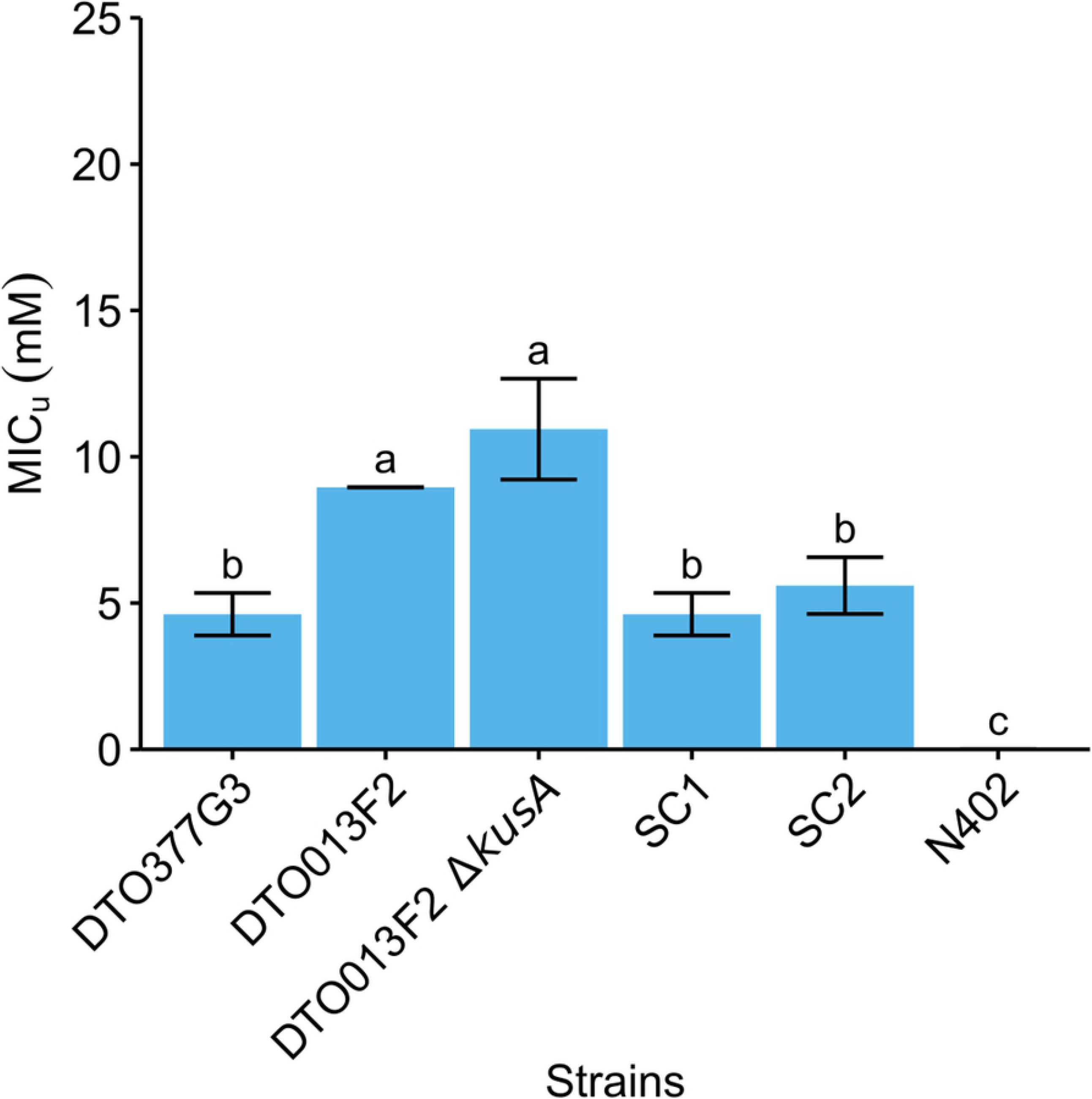
Sorbic acid resistance of five *P. roqueforti* strains and *A. niger* N402. The average MIC_**u**_ (mM ± standard deviation) is given for the fungal strains. Each bar graph represents average value of biological independent triplicates. Error bars indicate standard deviation and letters indicate significant difference in MIC_u_ (p < 0.05).

## Discussion

*P. roqueforti* is often encountered as a spoilage organism of food and feed. This can partly be attributed to its ability to grow at refrigeration temperatures [23], low O_2_ levels [24] and/or its resistance to preservatives, such as sorbic acid [25]. The inhibitory effect of propionic, benzoic and sorbic acid on *P. roqueforti* growth was assessed and benzoic acid was found to have the strongest inhibitory effect on 34 *P. roqueforti* strains. Previous studies report that *P. roqueforti* is resistant to benzoic acid. Growth was observed up to levels of 3000 ppm sodium benzoate [26,27], while we observed growth at 610 ppm (5 mM). Propionic acid hardly affected the growth of *P. roqueforti*, which is not surprising since it has already been reported that this fungus germinates on potato dextrose agar containing 0.5 M propionic acid (pH 5.6) with an estimated MIC of 0.79 M [23]. The sorbic acid resistance of *P. roqueforti* has also been assessed previously [14,25–30], reporting MIC values ranging from 0 – 40 mM sorbic acid (i.e. a MIC_u_ = 0 – 20 mM). This is similar to the range (MIC_u_ = 4.2 – 21.3 mM) found in this study. In fact, a resistant and sensitive group (R- and S-types) consisting of six and 28 strains, respectively, were found in this study. S-type and R-type strains showed a resistance up to 11.9 mM and 21.2 mM undissociated sorbic acid, respectively.

The R-type but not the S-type strains were found to contain a gene cluster (SORBUS) containing 70 genes, of which 51 genes are unique for the R-type strains. Even though the R-type strain DTO006G7 was used with the least fragmented Illumina assembly, the limits of the SORBUS cluster could not be established based on these results alone. The Oxford Nanopore of the DTO013F2 Δ*kusA* assembly revealed that SORBUS is part of a 2.3 Mb contig. Genes homologous to sorbic acid degradation-associated genes in *A. niger* (*sdrA, cdcA, padA, warA*) were identified in the *P. roqueforti* genome [9,12]. A total of 1, 4, 3 and 1 orthologs were found, respectively, and only *warA* and one *cdcA* ortholog were not located on the SORBUS cluster. Two genes with putative transmembrane transport (g216) or cation transporter (g296) function were identified in the PLINK analysis. The encoded proteins might also be involved in sorbic acid stress mediation alongside the SORBUS cluster, because in addition to decarboxylation, sorbic acid stress could be mediated by an efflux pump or through removal of protons from the plasma membrane by H^+^-ATPase [12,31].

As mentioned, only six out of the 34 strains assessed in this study were found to contain the SORBUS cluster. This might be explained by the ecology of *P. roqueforti. P. roqueforti* is found in forest soil and wood, but is also associated with lactic acid bacteria (e.g. in silage). Frisvad & Samson [32] already speculated that these micro-organisms may have very well co-evolved, because all the metabolics produced by lactic acid bacteria (e.g. lactic and acetic acid, CO_2_) are tolerated by *P. roqueforti* [33]. The elevated levels of weak acids might act as selection pressure to maintain SORBUS in *P. roqueforti* strains which grow in this niche environment. In contrast, this selection pressure is not present in cheese which might explain that none of *P. roqueforti* strains in the ‘cheese’ population contain the SORBUS cluster [18]. It should be noted that the S-type sequence fragments that align with SORBUS mostly consists of proteins annotated as transposase-like proteins or reverse transcriptase, which might explain why these fragments are found in the S-type strains and it suggests that SORBUS has been obtained from a different species by horizontal gene transfer. This is supported by the phylogenetic analysis on PFAM domains of five SORBUS genes, as the results show that three out of the five SORBUS specific genes are more closely aligned to *Aspergillus* species than *Penicillium* species. The gene cluster SORBUS was also present in two of 35 previously sequenced *P. roqueforti* strains [18] and their sorbic acid resistance could be determined to validate the role of SORBUS in sorbic acid resistance.

Transcriptome analysis of the R-type *P. roqueforti* strain DTO006G7 revealed that two of the three *cdcA* paralogs (*cdcA* and *cdcC*) are significantly down-regulated during growth in the presence of sorbic acid. This is in contrast with the results previously described [9], where the authors found a > 500-fold change for *cdcA* when *A. niger* was cultivated on sorbic acid. This difference might be caused by the difference in medium type, because that study [9] used sorbic acid as the sole carbon source, whereas in our experiments sorbic acid was used as a stressor in a nutrient-rich medium. This indicates that the sorbic acid content in the medium does not increase gene expression of the loci leading to increased resistance, suggesting that either these genes are constitutively expressed or expression is induced based on a different compound present in MEB.

Two partial SORBUS knockout strains in the DTO013F2 Δ*kusA* R-type strain showed reduced sorbic acid resistance to a level similar to that of the S-type strains. In contrast to the DTO013F2 Δ*kusA* strain, the SORBUS knockout strains were not repaired through homology directed repair using the donor DNA nor non-homologous end-joining. This might be due to the size of the fragment. Possibly, an alternative DNA repair mechanism such as microhomology-mediated end joining as described in *A. fumigatus* is employed in the DTO013F2 Δ*kusA* strain [34].

Despite the absence of a full SORBUS cluster in the S-type strains and the deletion strain, their MIC is still relatively high when compared to other fungal species such as *A. niger* (as shown in Fig 10) or *A. fumigatus* [12]. This suggests that along with the genes on the SORBUS cluster, other proteins are involved in sorbic acid stress mitigation. Our transcriptomics analysis revealed 21 down-regulated and 33 up-regulated genes that were similarly expressed both in a S-type and a R-type strain when exposed to sorbic acid. One of these up-regulated genes is the cation transporter (g1689, Table S4) which could have H^+^ ATPase activity in the plasma membrane to counteract the acidification caused by the undissociated sorbic acid in the cytosol [31].

In conclusion, the results presented in this study demonstrate that weak-acid resistance varies between *P. roqueforti* strains and the SORBUS cluster contributes to a high sorbic acid resistance. Yet, even in the absence of this cluster the resistance is still relatively high, implying that other mechanisms are also involved in resistance to this weak acid.

## Methods

### Strain and cultivation conditions

All fungal strains (Table S1) were provided by the Westerdijk Fungal Biodiversity Institute. Conidia were harvested with a cotton swab after seven days of growth at 25 °C on malt extract agar (MEA, Oxoid, Hampshire, UK) and suspended in 10 mL ice-cold ACES buffer (10 mM N(2-acetamido)-2-aminoethanesulfonic acid, 0.02 % Tween 80, pH 6.8). The conidia suspension was passed through a syringe containing sterilized glass wool and washed twice with ACES buffer after centrifugation at 4 °C for 5 min at 2,500 *g*. The spore suspension was set to 2·10^8^ spores mL^-1^ using a Bürker-Türk haemocytometer (VWR, Amsterdam, The Netherlands) and kept on ice until further use.

### Weak acid growth assay

Conidial suspension (5 µL) was inoculated in the centre of MEA plates containing 5 mM potassium sorbate, benzoic acid, or sodium propionate (all from Sigma). Medium was set at pH 4.0 using HCl, which corresponds to undissociated concentrations of 4.26, 3.07 and 4.42 mM of these acids, respectively. The absence of preservative was used as a control. Cultures were photographed after 5 days and colony surface area was measured using a manual threshold in ImageJ.

### Sorbic acid resistance

Conidial suspensions of the *P. roqueforti* strains were diluted to 10^7^ spores mL^-1^ and mixed in a 1:99 ratio with MEB pH 4.0 with and without 25 mM potassium sorbate. 300 µL of the resulting mixture was added in a well of a 96 wells plate (Greiner Bio-One, Cellstar 650180, www.gbo.com). Serial dilutions were made by mixing 225 µL MEB with potassium sorbate and 75 µL MEB without potassium sorbate, resulting in wells with potassium sorbate concentrations of 25, 18.75, 14.06, 10.55, 7.91, 5.93, 4.45 and 0 mM. This corresponded to undissociated sorbic acid concentrations of 21.22, 15.92, 11.94, 8.95, 6.72, 5.04, 3.78 and 0 mM, respectively. The undissociated sorbic acid concentrations were determined using the Henderson-Hasselbach equation.

The 96-wells plates were sealed with parafilm and incubated for 28 days at 25 °C using biologically independent replicates. After 28 days, growth was assessed and the undissociated minimal inhibitory concentration (MIC_u_) was determined for each strain. The MIC_u_ was defined as the lowest undissociated concentration in which no hyphal growth was observed. An one-way ANOVA followed by a Tukey’s HSD test was used to test for significant differences in MIC_u_ (*P* < 0.05).

### DNA extraction, genome sequencing, assembly and annotation

DNA extraction was performed as described [35] and Illumina NextSeq500 2×150 bp paired-end technology was used for sequencing (Utrecht Sequencing Facility, useq.nl). The reads were trimmed on both ends when quality was lower than 15 using bbduk from the BBMap tool suite (BBmap version 37.88; https://sourceforge.net/projects/bbmap/). The trimmed reads were assembled with SPAdes v3.11.1 applying kmer lengths of 21, 33, 55, 77, 99 and 127 and the –careful setting was used to reduce the number of indels and mismatches [36]. Genes were predicted with Augustus version 3.0.3 [37] using the parameter set that was previously generated for *P. roqueforti* [35]. Functional annotation of the predicted genes was performed as described [38]. Repetitive sequences in the assembly were masked using RepeatMaker [39], RepBase library [40] and RepeatScout [41]. The Short Read Archive (SRA) numbers of the datasets in this study are listed in Table S1 under accession numbers. The genomes, gene predictions and functional annotations can be accessed interactively at https://fungalgenomics.science.uu.nl. [Available upon publication].

### Genomic phylogeny and analysis

Single-copy orthologous groups were identified and aligned using OrthoFinder v2.5.2 [42]. A maximum likelihood (ML) inference was performed using RAxML [43]□ under the PROTGAMMAAUTO model. The number of bootstraps used was 200 (Average WRF = 0.43 %) and *Penicillium rubens* Wisconsin 54-1255 [44] was used to root the tree. The phylogenetic tree was visualized using iTOL v5 [45].

### Genome-wide association study

A genome-wide association study (GWAS) was performed based on the sorbic acid resistance screening and the whole-genome sequences. First, *P. roqueforti* strains were grouped into a resistant (R-type) or sensitive (S-type) group. DTO006G7 was selected from the R-group as reference for the analysis, because the assembly of this strain was the least fragmented in this group. Next, the program nucmer from the MUMmer suite (http://mummer.sourceforge.net/, version 4.0) was used to perform whole-genome alignment. Each genome was aligned to the reference and regions and genes unique for the R-type isolates were identified with the BEDtools package. The genome alignment was visualized with pyGenomeTracks [46]. A gene was considered absent when 90 % or more of its sequence was not found in the genome of a strain. The best practices recommended by GATK (Genome Analysis Toolkit) were used to obtain single nucleotide polymorphisms (SNPs) for each strain. In short, the sequence reads were aligned to the reference genome (DTO006G7) using Bowtie2 (version 2.2.9) and PCR duplicates were removed with Picard tools (MarkDuplicates; version 2.9.2). For variant calling, the HaplotypeCaller (GATK, version 3.7) was used with the following parameters: - stand_call_conf 30, -ploidy 1 and -ERC. The single-sample variant files (GVCFs) were joined into a GenomicsDB before joint genotyping. The variants were annotated using SNPeff (version 4.3) based on their predicted biological effect, such as the introduction of an early stop-codon or a synonymous annotation. The number of SNPs and their putative impact (‘low’, ‘moderate’ or ‘high’, as defined by SNPeff) was listed for each gene per genome. The SNPs were then correlated to sorbic acid resistance using PLINK v1.9 [21] with parameters ‘–maf 0.5 –allow-extra-chr’. The resulting association files were analysed and visualized using R.

### RNA extraction and sequencing

A genome-wide transcriptome analysis was performed on the sorbic acid sensitive *P. roqueforti* strain DTO377G3 and the sorbic acid-resistant *P. roqueforti* strain DTO006G7. Erlenmeyer flasks containing 50 mL MEB (pH 4.0) were inoculated with 100 µl ACES containing 10^7^ conidia and incubated for 48 h at 25 °C and 200 rpm. Mycelium was harvested using a sterilized miracloth filter and equally divided in Erlenmeyer flasks with 50 mL MEB (pH 4.0) or 50 mL MEB containing 3 mM potassium sorbate (pH 4.0). Growth was continued for another four hours, after which the mycelium was harvested using a sterilized Miracloth filter and frozen in liquid nitrogen. Total RNA was isolated with the RNeasy Plant Mini Kit (Qiagen) and purified by on-column DNase digestion according to the manufacturer’s protocol. RNA was sequenced with Illumina NextSeq2000 2×50 bp paired-end technology (Utrecht Sequencing Facility; useq.nl). The transfer experiment and subsequent RNA-sequencing was performed in biological triplicates. The transcript lengths, counts per gene and read mapping were determined using Salmon v1.5.2 with --validateMappings [47]. The transcript abundance of reads was quantified using custom constructed indices for DTO006G7 and DTO377G3.

DESeq2 [48] was used for pairwise comparisons of the samples and the identification of differentially expressed genes. Genes with low read counts (<10) were excluded from the analysis and a gene was considered differentially expressed when the adjusted p-value was < 0.05. In addition, genes were considered up- or down-regulated when they had a log_2_ fold change of > 2 or < −2, respectively. A Fisher Exact test as implemented in PyRanges [49] was employed to identify over- and under-representation of functional annotation terms in sets of genes. To correct for multiple testing the False Discovery Rate method was used, with a P-value < 0.05 as cut off.

The sequence reads are available in the Short Read Archive under BioProject PRJNA796729 [available upon publication].

### Plasmid construction and generation knockout strain

A *kusA* deficient DTO013F2 strain was constructed using the protocol and plasmid pPT22.4 as described [50]. Deletion was confirmed through diagnostic PCR and nanopore sequencing (Fig S1). CRISPR/Cas9 technology was employed to remove a 93 kbp region (from 84 to 177 kbp) of the SORBUS cluster in DTO013F2 Δ*kusA*. Two single-guide RNAs (sgRNAs) were designed to perform a simultaneous double restriction in the 93 kbp SORBUS region. Transformation procedures were performed as described, with some modifications [50]. In short, plasmid pFC332 [51] was used as a vector to express the sgRNA, *cas9* and a hygromycin selection marker (for primer sequences used in this study see Table S2). The 5’ and 3’ flanking regions of sgRNA were amplified using plasmids pTLL108.1 and pTLL109.2 as template [52]. The amplified products were fused and introduced into pFC332 using Gibson assembly (NEBuilder HiFi DNA Assembly Master Mix, New England Biolabs, MA, USA). The vectors containing the sgRNA were then transformed into competent *Escherichia coli* TOP10 cells for multiplication overnight. Plasmids were recovered using Quick Plasmid Miniprep Kit (ThermoFisher, Waltham, MA, USA) and digested using SacII to verify the presence of the sgRNA. In addition, the correct integration of the sgRNA was confirmed with sequencing. To construct donor DNA, two 1 kbp homologous regions located at scaffold 43 at nucleotide position 83573 to 84629 and 177760 to 178854 were amplified and fused using a unique 23 nucleotide sequence GGAGTGGTACCAATATAAGCCGG with a PAM site for further genetic engineering.

Transformation was performed as described with adjustments [53]. In short, *P. roqueforti* conidia were incubated 48 h at 25 °C in 100 mL potato dextrose broth at 200 rpm. The mycelium was washed in SMC and incubated for 4 h at 37 °C in lysing enzymes from *Trichoderma harzianum* (Sigma) dissolved in SMC. Protoplasts were resuspended in 1 mL STC and kept on ice after centrifuging for 5 min at 3000 g. To 100 μL of this suspension, 2 μg donor DNA, 2 μg of each pFC332 vector containing sgRNA (pTF and pSdrA) and 1.025 mL of freshly made PEG solution was added (see Table S3 for the vectors used in this study). After 5 min, 2 mL STC was added and the protoplasts were mixed with 20 mL liquid MMS containing 0.3 % agar and 200 μg hygromycin mL^-1^ (InvivoGen, San Diego, CA, USA). The mixture was poured on MMS containing 0.6 % agar and 200 μg hygromycin mL^-1^. Transformants were grown for 7-14 days at 25 °C and then single streaked on MM containing 100 μg mL^-1^ hygromycin until sporulating colonies appeared. Next, the plasmid was removed by a single streak on MM plates without antibiotic. Finally, transformants were single streaked on MM, MM containing 100 μg hygromycin mL^-1^ and MEA plates to confirm that transformants lost the plasmids.

To verify the transformants, genomic DNA from DTO013F2 Δ*kusA* and two DTO013F2 Δ*kusA* Δ*SORBUS* strains was sequenced with Oxford Nanopore MinION technology (FLO-MIN106) at the Utrecht Sequencing Facility (useq.nl). The reads were assembled using Canu v2.2 using the option for raw nanopore data and guided by a genome size of 28 Mbp [54]. In addition, the nanopore reads were aligned to the DTO006G7 assembly using Minimap2 [55]. Only reads aligning once and that had a mapping quality > 60 were selected using samtools. The sequencing data generated with the MinION technology is deposited in the SRA archive under the following accession numbers: SRR17178875, SRR17178876 and SRR17178877.

## Acknowledgements

We thank Utrecht Sequencing Facility (useq.nl) for providing sequencing service and data. Utrecht Sequencing Facility is subsidized by the University Medical Center Utrecht, Hubrecht Institute, Utrecht University and The Netherlands X-omics Initiative (NWO project 184.034.019).

